# Consequences of light spectra for pigment composition and gene expression in the cryptophyte *Rhodomonas salina*

**DOI:** 10.1101/2023.09.20.558689

**Authors:** Rachel A. Schomaker, Tammi L. Richardson, Jeffry L. Dudycha

**Author notes:** Author contributions: JLD and TLR conceived the work, RAS generated and analyzed data, RAS, TLR, and JLD wrote the manuscript.

## Abstract

Algae with a more diverse suite of pigments can, in principle, exploit a broader swath of the light spectrum through chromatic acclimation, the ability to maximize light capture via plasticity of pigment composition. We grew *Rhodomonas salina* in wide-spectrum, red, green, and blue environments and measured how pigment composition differed. We also measured expression of key light-capture and photosynthesis-related genes and performed a transcriptome- wide expression analysis. We observed the highest concentration of phycoerythrin in green light, consistent with chromatic acclimation. Other pigments showed trends inconsistent with chromatic acclimation, possibly due to feedback loops among pigments or high-energy light acclimation. Expression of some photosynthesis-related genes was sensitive to spectrum, although expression of most was not. The phycoerythrin α-subunit was expressed two-orders of magnitude greater than the β-subunit even though the peptides are needed in an equimolar ratio. Expression of genes related to chlorophyll-binding and phycoerythrin concentration were correlated, indicating a potential synthesis relationship. Pigment concentrations and expression of related genes were generally uncorrelated, implying post-transcriptional regulation of pigments. Overall, most differentially expressed genes were not related to photosynthesis; thus, examining associations between light spectrum and other organismal functions, including sexual reproduction and glycolysis, may be important.

**Originality-Significance Statement:** Most work on light and algal photophysiology focuses on light intensity rather than light spectrum. Given the large spectral variation of light in aquatic systems, explaining how such algae respond to spectral variation will provide a better foundation for understanding the base of aquatic food webs. Much of the light spectrum is poorly absorbed by chlorophyll, which creates an opportunity for photosynthetic species with other pigments. We quantified physiological and genetic responses to light spectrum in replicate experimental populations of *Rhodomonas salina*, an alga with a phycoerythrin in addition to chlorophylls. We predicted photophysiology and gene expression would change to maximize *R. salina’s* capacity to capture available light, in accordance with the theory of chromatic acclimation. Our results show that responses to light spectra are more complex than predicted. Some aspects of photophysiology did support the theory’s predictions, but gene expression was generally unrelated to variation of light spectrum or photophysiology. This not only suggests that chromatic acclimation is potentially regulated post-transcriptionally, but also that physiological processes – notably glycolysis and the transition to sexual reproduction – that may be regulated by light spectrum. Our work adds to the generally limited work on light spectrum and physiology by investigating a eukaryote from a phylum with a great diversity of photosynthetic pigments.

## Introduction

Photosynthesis is the remarkable metabolic process whereby organisms capture light energy and use it to fix carbon dioxide into organic carbon compounds. The evolution of photosynthesis resulted in an explosion of biodiversity across the globe, and understanding the functionality, plasticity, and ecological consequences of this process remains an area of substantial interest.

Modern photosynthetic organisms have evolved to use chlorophyll-*a* in their reaction centers to funnel light energy through the photosynthetic pathway. Chlorophyll-*a* is excellent at absorbing blue (∼400 – 490 nm) and red (∼620 – 700 nm) wavelengths of the visible spectrum but it does not efficiently absorb the remaining wavelengths, leaving a wide range of potentially untapped energy that could be used for photosynthesis (Mackinney 1941).

Accessory pigments are light-absorbing compounds that differ among algal taxa and work in conjunction with chlorophyll-*a* by capturing light that chlorophyll-*a* absorbs poorly (Blinks 1954; Glazer 1977; Gantt 1980; Stengel *et al*. 2011). As a result, phytoplankton accessory pigments may open spectral niches that were not previously available, which can lead to increased biodiversity within ecosystems, altered community dynamics, and ecosystem functioning (Stengel *et al*. 2011; Sanfilippo *et al*. 2019). These potential effects of niche differentiation and exploitation can have substantial downstream effects at both the community and ecosystem level, especially because the spectral characteristics of aquatic environments can vary substantially in time and space. Aquatic habitats rich in colored dissolved organic material (CDOM) tend to be dominated by red light because CDOM strongly absorbs blue and violet light (Blough and Del Vecchio 2002). Offshore oceans tend to be dominated by blue light because they have low CDOM and low phytoplankton concentrations (Kirk 1994; Blough & Del Vecchio 2002), while coastal oceans are often green in appearance, as nutrient inputs promote phytoplankton growth and hence high chlorophyll-*a* concentrations. As anthropogenic land-use, eutrophication, and CDOM input into aquatic environments rise, the amount of spectral variation across aquatic habitats may become more extreme (Roulet & Moore 2006; Kritzberg 2017; Dutkiewicz *et al*. 2019; Luimstra *et al*. 2020). These changes could force natural phytoplankton populations into new spectral environments, which can alter the ecosystem if the resident organisms are unable to effectively occupy them.

One way phytoplankton can respond to shifts in the spectral environment is by adjusting the ratio of various pigments in response to the spectral environment (Sanfilippo *et al*. 2019; Sebelik *et al*. 2020). Known as chromatic acclimation (or chromatic “adaptation”), this is a form of reversible phenotypic plasticity where photophysiology is adjusted to maximize light absorption (Engelmann 1883; Gaidukov 1903; Hattori & Fujita 1959a,b; Fujita & Hattori, 1960a,b, 1962a,b, 1963; Bennett & Bogorad 1973). Chromatic acclimation is well-studied in cyanobacteria because many species maintain a diverse pigment complement, including the accessory pigments phycocyanin and phycoerythrin, which primarily absorb red (569-650nm) and green light (538- 568nm) respectively, along with chlorophylls (Campbell 1996; Stengel *et al*. 2011; Xia *et al*. 2016). Cyanobacteria shift their pigment composition and adjust the size of their phycobilisomes to best suit the light characteristics of their habitats, broadening their fundamental niche and giving them a competitive advantage where spectral variation occurs (Grossman 2003; Stomp *et al*. 2004; Stomp *et al*. 2007; Montgomery 2017). Beyond cyanobacteria, physiological responses to light spectrum are widespread across many different eukaryotic phytoplankton (Wallen & Geen 1971; Rivkin 1989; Algarra et al. 1991; Figueroa et al. 1995; Granbom et al. 2001; Mouget et al. 2004; Vadiveloo et al. 2017), and light spectrum has been shown to affect not only pigment composition in eukaryotic phytoplankton, but growth rate, as well (Heidenriech & Richardson 2019), suggesting that light spectrum has major implications for phytoplankton fitness. However, the molecular mechanisms of these responses are poorly known and research on how gene expression in algae responds to changes in spectral irradiance is largely lacking; most such work in algae involves the effects of light intensity but not light spectrum (e.g., Ho et al. 2009; Park et al. 2010; Xiang et al. 2015; Nan et al. 2018; Li et al. 2019). Our aim is to begin remedying this gap for a eukaryotic alga that has multiple photosynthetic pigments.Cryptophytes are a phylum of single-celled eukaryotic algae that are ubiquitous across nearly all aquatic habitats and exhibit remarkable diversity in visible pigmentation. Photosynthetic cryptophytes contain chlorophyll-*a* and also maintain the accessory pigments chlorophyll-*c_2_*, alloxanthin, α-carotene, and phycobiliproteins (cryptophyte phycoerythrin and cryptophyte phycocyanin). Unlike cyanobacteria, cryptophyte species each have only one type of phycobiliprotein (appearing either green or red), which are the main light-harvesting pigments in cryptophytes (Glazer 1983; Hill & Rowan 1989; Vesk *et al*. 1992; Blankenship 2002). These pigments are composed of two alpha and β protein subunits, plus four chromophores known as phycobilins. The molecular structure of the protein-chromophore complex is directly related to the wavelengths of light the pigment can capture (Doust *et al*. 2004; Overkamp *et al*. 2014).

Phycobiliprotein evolution is associated with changes in light capture in cryptophytes (Greenwold *et al*. 2019), but studies examining chromatic acclimation are conflicting or differ greatly among clades (Ojala 1993; Kamiya & Miyachi 1984 a,b; Lawrenz & Richardson 2017; Heidenreich & Richardson 2017). Our study aims to build off these previous observations in order to better understand cryptophytes’ ability to respond to light spectrum and to investigate the molecular responses, which have not been studied previously. To do this, we investigated the plasticity of pigment composition and gene expression (i.e., whether pigment composition and expression level changes) in the cryptophyte *Rhodomonas salina* (which has cryptophyte pycoerythrin-545) grown in different spectral environments. We asked the following questions: 1) How do the concentrations of pigments change in response to different spectral irradiance but equal intensity? and 2) How does gene expression differ among spectral environments? We examined the pigment and transcriptional responses for experimental populations grown in blue, red, and green spectra. The blue vs. red light comparison represents the widest energy difference between spectra (as one blue photon is more energetic than one red photon) and reflects distinct habitats in the natural world. Green vs. red light also reflects distinct real-world habitats but maximizes the differences in the expected light absorption due to molecular physiology. In contrast, blue vs. green light comparisons represent distinct habitats that present more limited energetic and light absorption differences. We also collected data for the wide-spectrum environment our culture of *R. salina* had been growing in prior to the experiment as a baseline. Based on the theory of chromatic acclimation, we expected that *R. salina* would respond to maximize its capacity to capture available light; if *R. salina* was not plasticly responsive to light color, we expected to see no change in *R.* salina’s physiology or gene expression across light spectra.

We investigated transcriptional responses at three different scales. First, we examined expression of transcripts that encode the peptide components of cryptophyte phycobiliproteins, predicting that they would correlate with concentration of the pigment and maximize available light capture. For example, we expected to see an increase in phycoerythrin concentration in green light, and we expected to see the genes encoding for the phycoerythrin subunit proteins to be upregulated in green light to mirror this shift in concentration. Assuming changes in concentration maximized light capture, we expected changes in concentration and gene expression to remain stable in order to maintain this ability. Second, we examined expression of 99 genes that were *a priori* identified as participating in light capture or photosynthesis, predicting that these loci would be most sensitive to light spectrum, but the direction of regulation would be dependent upon each gene’s function. Third, we examined genome-wide expression to identify molecular processes that interact with light spectrum but may not have obvious connections to light capture or photosynthesis. Because we do not know the specific function of every gene in the *R. salina* genome, we did not have any specific hypotheses for the direction of regulation we expected for many genes. We further sought to link our assessment of pigment plasticity (change in pigment composition) and expression plasticity (change in expression level) by testing for correlations between the two across our entire experiment.

## Results

### Pigment Data

#### Total pigments

Overall, there were no significant differences in total pigment across spectra. Concentrations were highest in green light (10.80 ± 1.72 pg/cell) followed by wide-spectrum cells, (7.40 ± 2.34 pg/cell), then blue (7.40 ± 2.80 pg/cell), and red light (7.10 ± 3.87 pg/cell). In all spectral environments, cryptophyte phycoerythrin comprised the greatest percentage of the total pigment concentration, followed by chlorophyll-*a*, chlorophyll-*c_2_*, alloxanthin, and α-carotene.

#### Phycoerythrin concentrations

We saw the highest cryptophyte phycoerythrin concentrations in cultures grown under green light (6.2 ± 0.60 pg/cell) and the lowest in those grown in red light (2.7 ± 0.60 pg/cell) (Figure 1a). Populations grown in blue light and the wide-spectrum light had average cryptophyte phycoerythrin concentrations of 4.6 ± 0.60 pg/cell and 3.8 ± 0.90 pg/cell, respectively (Figure 1a). The cryptophyte phycoerythrin concentrations were significantly different between populations grown in green and red light (*p*-value = 0.0069; *F*- value = 7.85; *df* = 2). We did not observe any other significant differences between the blue vs. red or blue vs. green comparisons. For wide-spectrum, blue, and green light environments, cryptophyte phycoerythrin comprised 50% or more of the total cellular pigment concentration (50.5%, 62.0%, and 57.4% for wide, blue, and green light, respectively). Cryptophyte phycoerythrin concentrations comprised only 37% of the total cellular pigment for cultures grown in red light (Figure 2).

**Figure 1:**
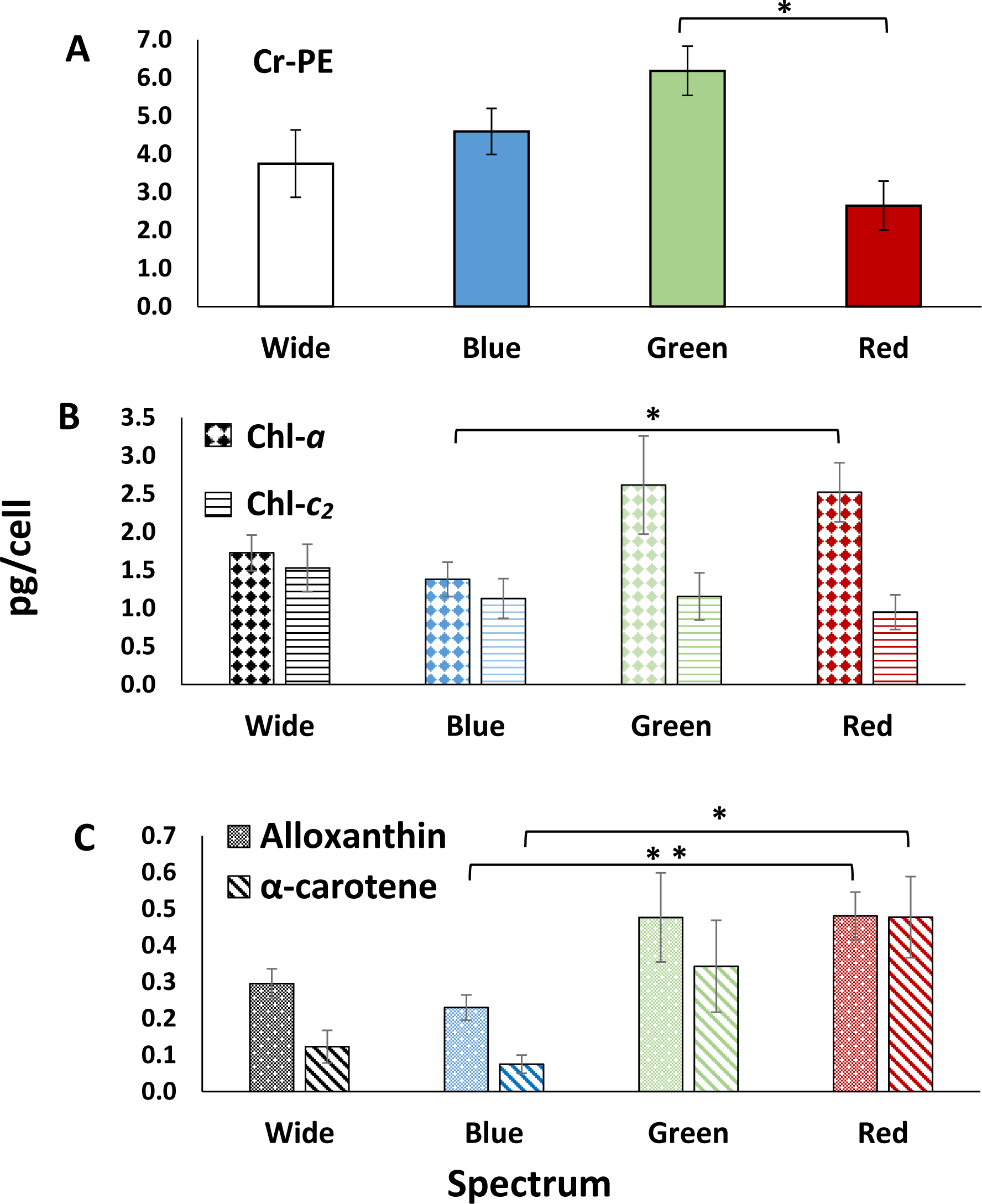
Average concentrations in pg/cell of all pigments in *R. salina* grown in wide-spectrum, blue, green, and red spectral environments. Error bars are standard error. Note the differences in the y-axis scale for all three graphs. **A)** Cryptophyte phycoerythrin (Cr-PE). **B)** Chlorophyll-*a* (chl-*a)* and chlorophyll-*c_2_* (chl-*c_2_*). **C)** Alloxanthin and α-carotene.

**Figure 2:**
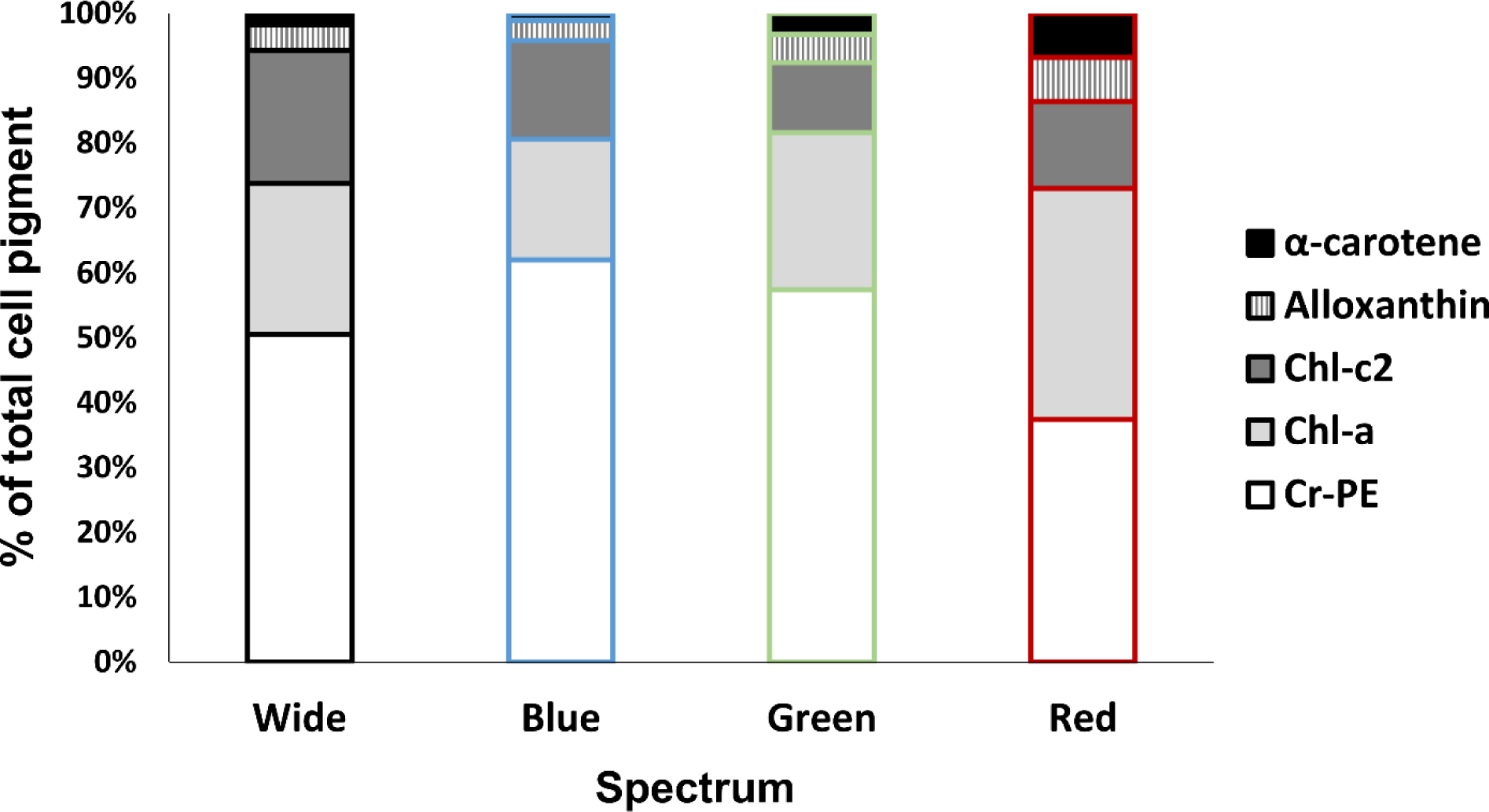
Relative pigment concentration of α -carotene, alloxanthin, chlorophyll-*c_2_* (chl-*c_2_*), chlorophyll-*a* (chl-*a*), and cryptophyte phycoerthrin (Cr-PE) across all four light treatments. Relative pigment concentrations were calculated on a mass/cell basis.

#### Non-PBP pigment concentrations

Chlorophyll-*a* concentrations were significantly different between populations grown in red (2.5 ± 0.40 pg/cell) and blue light (1.4 ± 0.20 pg/cell) (*p*-value = 0.045; Z = -2.00). Populations grown in green light had an average chlorophyll-*a* concentration of 2.6 ± 0.60 pg/cell, while those grown in the wide-spectrum environment had 1.7 ± 0.20 pg/cell (Figure 1b). While populations grown in green light exhibited the highest average chlorophyll-*a* concentrations compared to the other spectral habitats, there were no significant differences observed between green light chlorophyll-*a* concentrations and the other spectra. The percentage of total cellular pigment that is chlorophyll- *a* differed with environment, with wide-spectrum, blue, green, and red-light environments exhibiting chlorophyll-*a* percentages of 23.3, 18.6, 24.3, and 35.6%, respectively. Chlorophyll-*c_2_* concentrations differed slightly across the different spectral environments, but there were no significant differences between any spectral comparisons (Figure 2).

Alloxanthin and α-carotene concentrations (Figure 1c) were both significantly different between and red and blue treatments (alloxanthin *p*-value = 0.037; Z = -2.09; α-carotene *p*-value = 0.0061; Z = -2.74). Alloxanthin concentrations ranged from 4.0 ± 0.04, 3.1 ± 0.03, 4.4 ± 0.12, and 6.8 ± 0.07 pg/cell in wide-spectrum, blue, green, and red light, respectively, while α-carotene concentrations were 1.7 ± 0.04, 1.0 ± 0.02, 3.2 ± 0.13, and 6.7 ± 0.11 pg/cell.

### Transcriptome Assembly

Our final transcriptome assembly contained 24,167 contigs, had an N50 of 2,431 bp, and had a GC content of 58.75%. Publicly available cryptophyte transcriptome assemblies of species from the *Hemiselmis, Proteomonas, Cryptomonas, Chroomonas, Guillardia,* and *Rhodomonas* clades (Marine Microbial Eukaryote Transcriptome Sequencing Project) range from 24,119 to 41,208 contigs with varying assembly metrics. Our *R. salina* assembly statistics thus fall within the expected published range of cryptophyte transcriptomes. A previously assembled *R. salina* transcriptome based on 50 bp reads assembled with Abyss (MMETSP1047-20130122) had 31,523 contigs and an N50 of only 1,650. Any disparities observed are likely due to differences in assembly methods, sequencing depth, read length, or species’ biological variation.

Of the 24,167 contigs in our assembly, we were able to identify 13,170 (54.50%) transcripts by matching them to protein annotations from the Pfam-A, Rfam, OrthoDB, and uniref90 databases. The remaining transcripts did not return an annotation hit across the protein databases.

The results of our BUSCO analysis revealed that 63.0% of expected eukaryotic orthologs (determined by the number of total complete BUSCOs, both single copy and duplicated) are present in our assembly, while 32.7% and 46.5% of the complete chlorophyte and protist BUSCOs were present, respectively (Table 1) (Simao *et al*. 2015).

**Table 1:** BUSCO results breakdown of the completed *R. salina* assembly against the eukaryote, chlorophyte, and protist databases. C = Complete; S = Complete and single-copy; D = Complete and duplicated; F = Fragmented; M = Missing.

| BUSCO Category | Eukaryote Database | Chlorophyte Database | Protist Database |
| --- | --- | --- | --- |
| Complete BUSCOs (C)<br>(includes single-copy and duplicated) | 191 (63.0%) | 710 (32.7%) | 100 (46.5%) |
| Complete and single-copy BUSCOs (S) | 158 | 573 | 86 |
| Completed and duplicated BUSCOs(D) | 33 | 137 | 14 |
| Fragmented BUSCOs (F) | 21 (6.9%) | 92 (4.3%) | 0 (0%) |
| Missing BUSCOs (M) | 91 (30.1%) | 1366 (63.0%) | 115 (53.5%) |
| Total BUSCO groups searched | 303 | 2168 | 215 |

### Differential Gene Expression Analysis and Exploration

We found that neither of the cryptophyte phycoerythrin subunit genes were significantly differentially expressed in any of our three spectral comparisons (FDR *p*-value < 0.05 and a log_2_ foldchange ≥ 2). To evaluate the potential for false negatives, we relaxed the FDR and foldchange; this had no effect on the outcome in the blue vs. red or blue vs. green comparisons. However, when the FDR cutoff was adjusted to 0.1 with a foldchange of 1.5, then expression of the cryptophyte phycoerythrin β subunit gene was significantly downregulated in green light compared to red light – opposite the pattern of the pigment’s concentration. The cryptophyte phycoerythrin α and β subunit gene expression patterns did not match that of the cryptophyte phycoerythrin concentrations. For the α-subunit gene(s), we observed the highest expression in wide-spectrum light (2,254 TPM) and the lowest in blue (1,306 TPM) and red light (1,301 TPM), but for the β subunit, red light cultures exhibited the highest expression (103 TPM), while the wide-spectrum, blue, and green light cultures were nearly the same (72, 71, and 65 TPM, respectively) (Figure 3).

**Figure 3:**
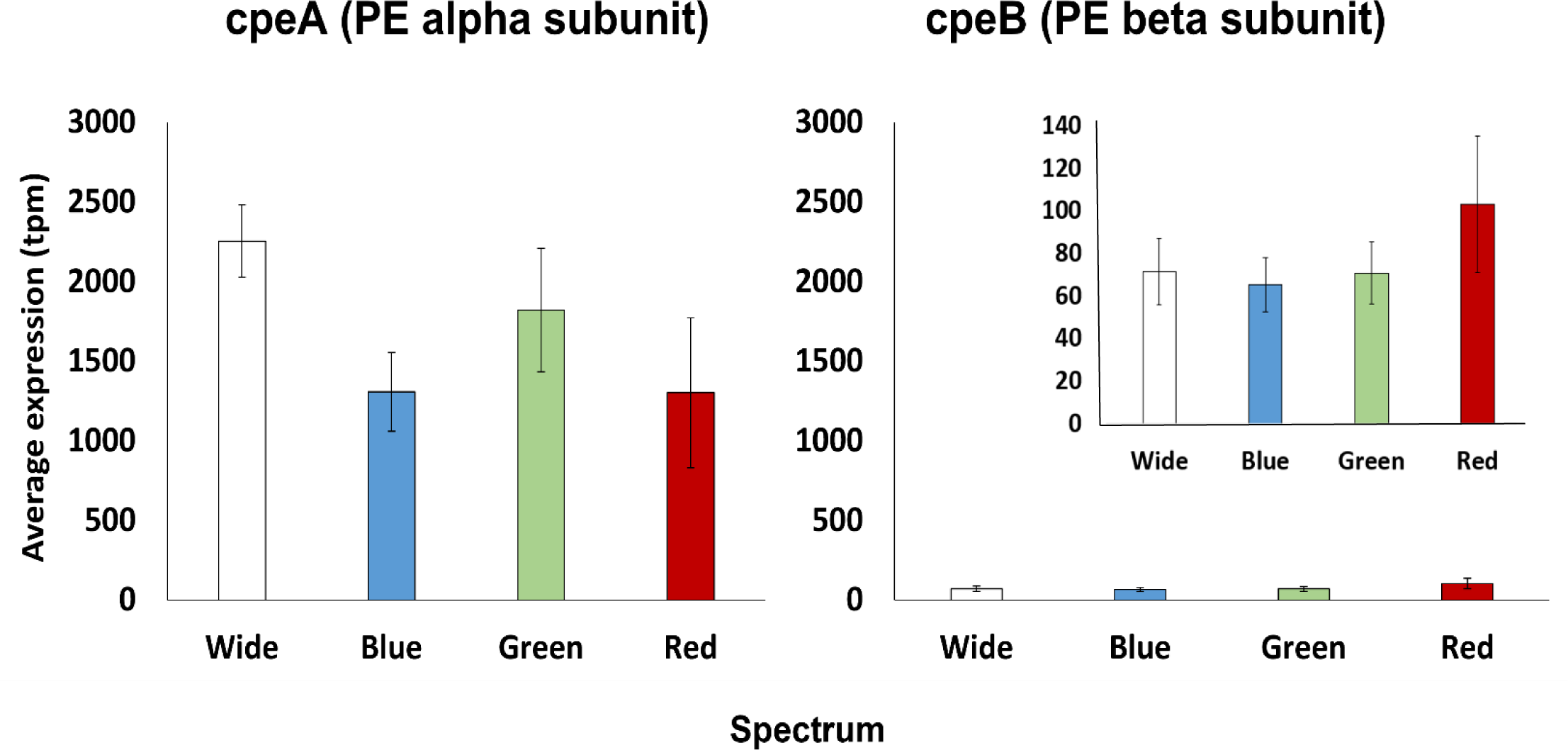
Expression of Cryptophyte phycoerythrin α and β subunits across the four different light environments. Expression values are reported in transcripts per million (tpm). There were no significant differences for either subunit across spectra. Note the difference in scales for the cpeB expression inset. Error bars represent standard error.

When we performed these same comparisons for our *a priori* photosynthesis gene set, we found that very few photosynthesis-related genes were significantly differentially expressed between light spectra. With the standard FDR of 0.05, 9 genes were differentially expressed in the blue vs. red treatments, while there were none in either the green vs. red or green vs. blue comparisons. When we relaxed the FDR to 0.1 with a log_2_ foldchange ≥ 2, we saw 12 differentially expressed genes in blue vs. red, 5 for green vs. blue, and still none for green vs. red. We did not see any evidence for differential expression of photosynthetic genes in the green vs. red comparison until we reached an FDR of 0.14 with no specified log_2_ foldchange, where we then had 2 genes returned. The top annotated differentially expressed genes for each of our photosynthetic comparisons are outlined in Table 2A-C.

**Table 2:** Significantly differentially expressed genes from the photosynthetic gene set for **A:** Blue vs Red; **B:** Blue vs Green; and **C:** Green vs Red comparisons.

**A: Blue vs Red – FDR of 0.1; no specified log<sub>2</sub> foldchange (i.e. log<sub>2</sub> foldchange ≥ 0)**
| Transcript ID from Assembly | Annotation | FDR p-value | Observed fold-change | Expression Direction |
| --- | --- | --- | --- | --- |
| Transcript_22761 | Chlorophyll binding protein | 0.0000526 | 2.76 | Up in Blue |
| Transcript_16447 | RUBISCO Large Subunit | 0.000221 | 2.64 | Up in Red |
| Transcript_23199 | ATP synthase CF1 alpha subunit | 0.0.00141 | 4.81 | Up in Red |
| Transcript_16630 | Photosystem I P700 chlorophyll a apoprotein A1 | 0.00123 | 2.82 | Up in Red |
| Transcript_21753 | Potential phycoerythrin α-subunit | 0.00109 | 2.84 | Up in Blue |
| Transcript_5484 | Photosystem II protein Y | 0.00269 | 1.40 | Up in Blue |
| Transcript_4751 | Chlorophyll binding protein | 0.00326 | 1.13 | Up in Red |
| Transcript_8725 | PSII Factor/Component | 0.00299 | 1.77 | Up in Red |
| Transcript_2439 | Chlorophyll binding protein | 0.00432 | 1.38 | Up in Red |
| Transcript_8558 | PSII Factor/Component | 0.00530 | 1.11 | Up in Red |
| Transcript_9593 | Photosystem II protein W | 0.01 | 1.51 | Up in Blue |
| Transcript_2914 | Potential phycoerythrin α-subunit | 0.01 | 0.95 | Up in Red |

**B: Blue vs Green - FDR of 0.1; no specified log<sub>2</sub> foldchange (i.e. log<sub>2</sub> foldchange ≥ 0)**
| Transcript ID from Assembly | Annotation | FDR p-value | Observed fold-change | Expression Direction |
| --- | --- | --- | --- | --- |
| Transcript_2914 | Potential α-phycoerythrin subunit | 0.00316 | 1.12 | Up in Green |
| Transcript_21753 | Potential α-phycoerythrin subunit | 0.00224 | 2.50 | Up in Blue |
| Transcript_2439 | Chlorophyll binding protein | 0.00140 | 1.56 | Up in Green |
| Transcript_22761 | Chlorophyll binding protein | 0.00153 | 2.02 | Up in Blue |
| Transcript_4815 | Phycoerythrin lyase | 0.00345 | 1.19 | Up in Blue |

**C: Green vs Red - FDR of 0.14; no specified log<sub>2</sub> foldchange (i.e. log<sub>2</sub> foldchange ≥ 0)**
| Transcript ID from Assembly | Annotation | FDR p-value | Observed fold-change | Expression Direction |
| --- | --- | --- | --- | --- |
| Transcript_23199 | ATP synthase CF1 alpha subunit | 0.00276 | 4.38 | Up in Red |
| Transcript_4815 | Phycoerythrin lyase | 0.00212 | 1.26 | Up in Red |

When we ran the analysis for the complete gene set, we found that 1,290 genes were significantly differentially expressed in the blue vs. red comparison with an FDR *p*-value < 0.05 and a log_2_ foldchange ≥ 2. Of these, 990 were upregulated in blue light, while 300 were upregulated in red light (Figure 4; Figure 5; Figure S2a). Our gene ontology results suggest that the genes upregulated in blue light were involved in a wide array of functions, including oxidation-reduction processes, translation, transmembrane transport, carbohydrate metabolic processes, transcriptional regulation, cell signaling, transduction, and communication, and various biosynthetic processes, such as phospholipid and nucleotide biosynthesis. Those upregulated in red light were primarily involved in translation, transmembrane proteins and ion transport, oxidation-reduction processes, and photosynthetic electron transport.

**Figure 4:**
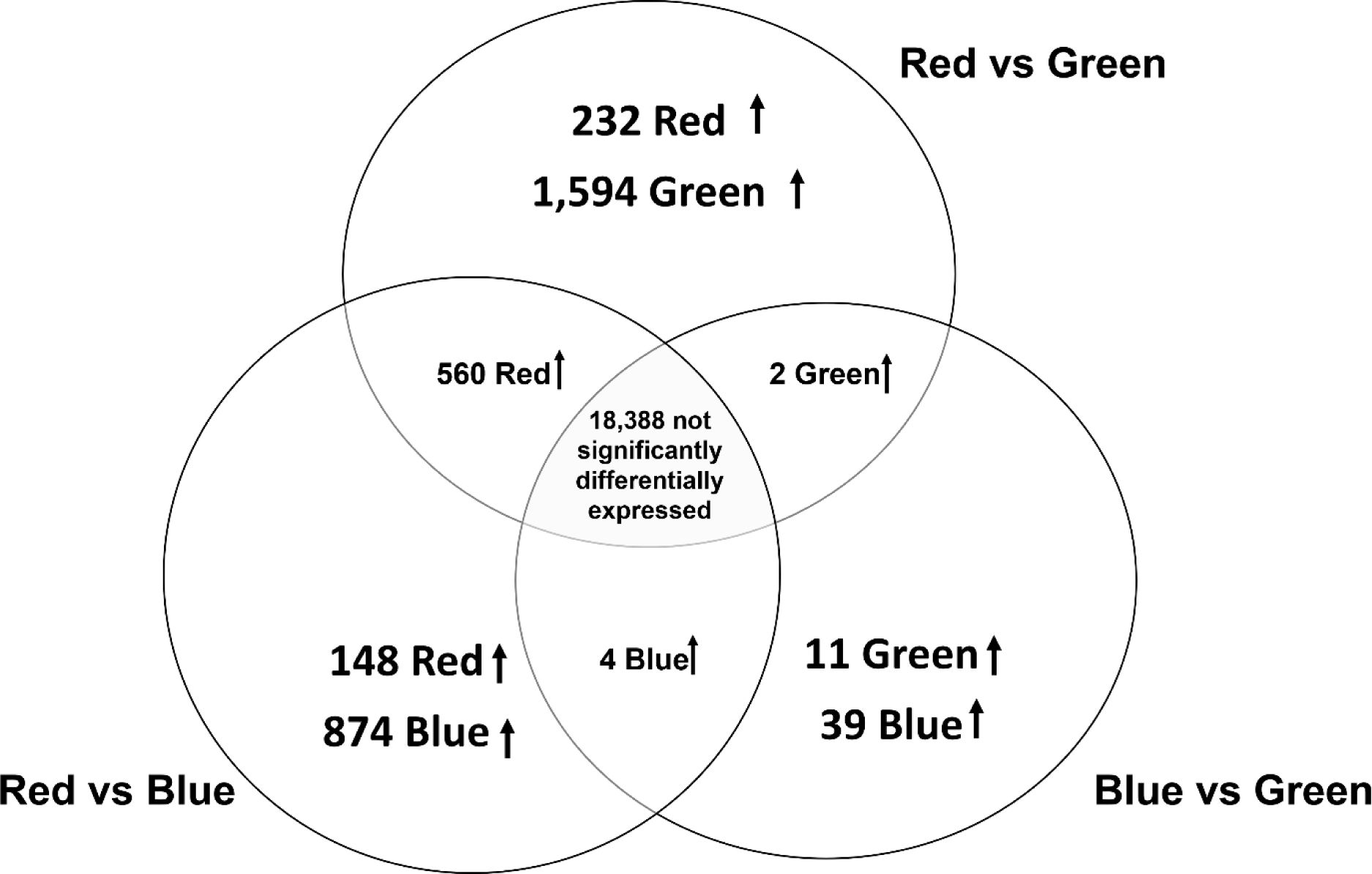
Venn diagrams for all comparisons. Numbers shown in each comparison circle represent genes upregulated in each spectral environment compared to the other. Numbers displayed between comparisons represent genes upregulated in the given spectral environment regardless of which light environment was the opposing comparison. The value in the center represents the number of genes that were not significantly differentially expressed in any comparison.

**Figure 5.**
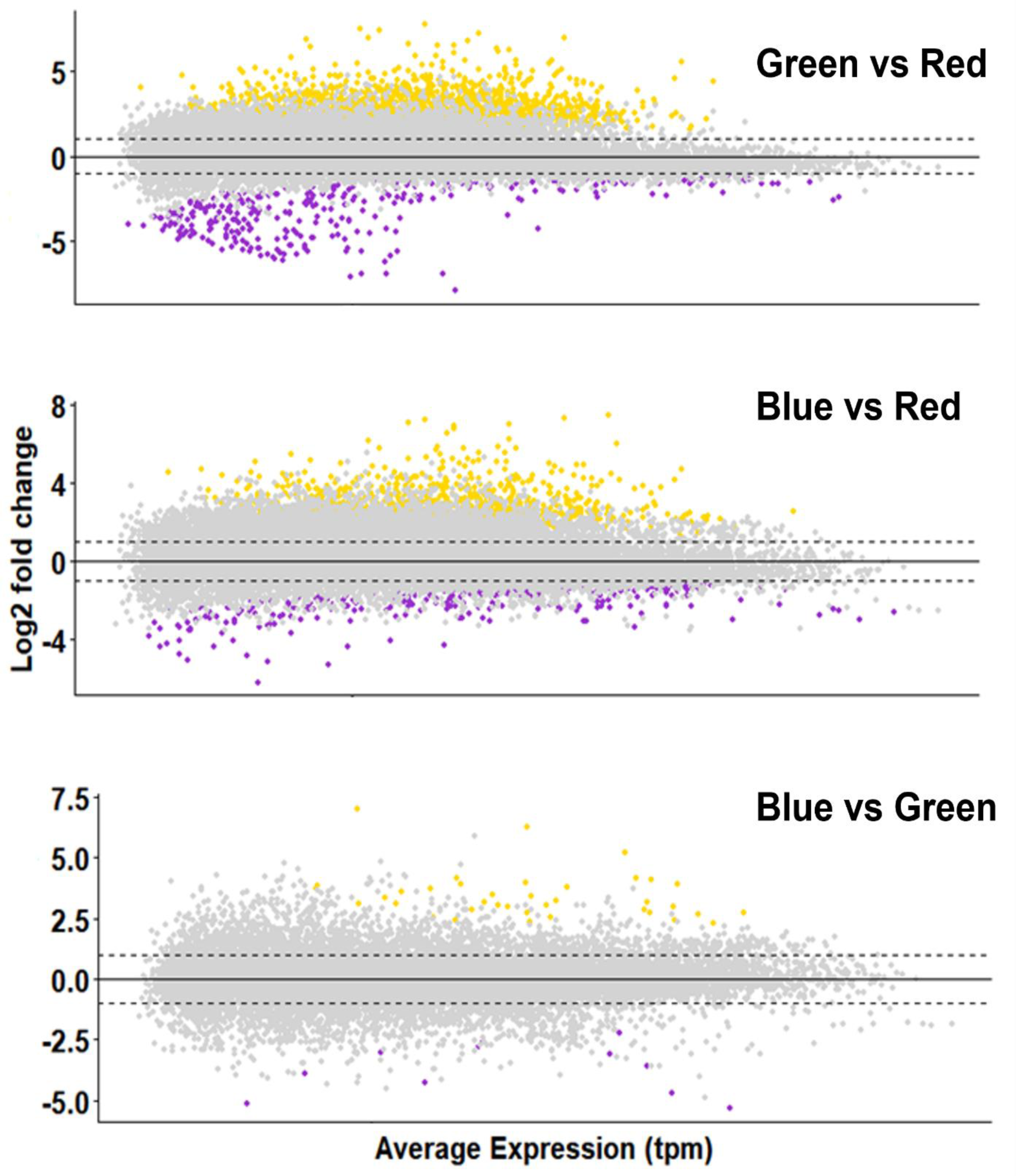
MA plots for all comparisons. For Green vs Red, differentially expressed genes relative to expression in red light. For Blue vs Red, differentially expressed genes are relative to expression in red light. For Blue vs Green, differentially expressed genes are relative to green light. Yellow dots represent significantly upregulated genes; purple dots represent significantly downregulated genes. Grey dots represent expressed genes which were not significant. The FDR *p*-value cutoff was 0.05 with a log_2_ fold-change > ±2 (represented by the dashed horizontal lines).

For the green vs. red comparison, 1,826 genes were significantly differentially expressed (FDR p-value < 0.05; log_2_ foldchange ≥ 2) (Figure 4; Figure 5; Figure S2b). Of these 1,826, only 232 were upregulated in red light and 1,594 were upregulated in green light. Both green- and red-light environments saw an upregulation of different genes involved in biological processes primarily involved in oxidation-reduction processes, transmembrane transport, and protein phosphorylation. Green light also upregulated genes involved in carbohydrate metabolic processes, transcription and translational regulation, DNA replication and repair, and multiple RNA processing mechanisms, including mRNA, rRNA, and tRNA processing, mRNA splicing, RNA polymerase regulation, and mRNA catabolism.

Only fifty genes were significantly differentially expressed between blue and green light.

Of these, 39 were upregulated in blue light, while 11 were upregulated in green light (FDR p-value < 0.05; log_2_ foldchange ≥ 2) (Figure 4; Figure 5; Figure S2c). The fifty genes included ones that were functionally involved in methyltransferase activity, tyrosine phosphatase function, DNA binding, transcriptional regulation, and protein folding and transport.

Many of the top differentially expressed genes could not be identified across any of the protein databases used in our dammit annotation pipeline, nor when we tried identifying their potential annotation using the NCBI conserved domain database (CDD) (Lu *et al*. 2020). Thus, it is apparent that some transcriptionally active and spectrally responsive regions of the *R. salina* genome are currently unannotated and may require extension or deeper exploration.

### Correlation Analysis

We found no significant correlations between cryptophyte phycoerythrin pigment concentration and protein subunit expression (both α and β subunits). When we tested for correlations among expression patterns of the photosynthetic gene set and the various pigment concentrations, we found a total of 14 genes were significantly (*p*-value < 0.05) correlated with chlorophyll-*a* and chlorophyll-*c_2_* concentration (Supplemental Table 2). Most of these genes were related to PSI and PSII synthesis, chlorophyll binding proteins, or ATP synthesis, and two were related to cryptophyte phycoerythrin synthesis or function. One gene encoding for a chlorophyll A-B binding protein (transcript 22761 in our assembly), a protein that binds to chlorophyll to form light-harvesting complexes (Dittami *et al*. 2010; Sturm *et al*. 2013; Hey and Grimm 2020), was significantly correlated with cryptophyte phycoerythrin concentration (coefficient = 0.499, *p*-value = 0.049). Twenty-four genes were significantly correlated with the photoprotective pigments (Supplemental Table 3). Of these twenty-four, six were chlorophyll binding proteins that all exhibited negative correlations; five were related to ATP synthesis that all had positive correlations; nine were proteins for PSI or PSII synthesis and function, which were mostly positively correlated; and one was the RuBisCo large subunit (rbcL), which had a significantly positive correlation.

## Discussion

For aquatic photosynthetic organisms, light is often a major limiting resource and cause of competition, thus organisms efficient at exploiting a broader range of light colors should have an advantage over those with more limited absorption options. We examined the physiological plasticity in pigment composition and gene expression in *R. salina* grown in wide-spectrum, blue, green, and red light. We expected that differences in pigmentation would follow the general theory of chromatic acclimation (Engelmann 1883, 1902; Gaiducov 1902, 1903), where pigments (type and concentrations) adjust to optimize absorption of wavelengths of available light, and we expected gene expression of pigment-related genes to follow the same pattern. We also predicted that other photosynthesis-related genes would have high expression sensitivity to changes in light color, but we did not expect that many non-photosynthesis-related genes would respond to spectrum. Our data partially supported the theory of chromatic acclimation, but some deviations from expected pigment concentrations and the lack of clear drivers at the level of gene expression suggest that plastic responses to light spectrum are more complex than the theory assumes.

### Are cryptophyte pigments maximizing their capacity to capture available light?

*R. salina* contains cryptophyte phycoerythrin 545, which efficiently absorbs green wavelengths of light (with a maximum absorption peak at 545 nm). Because of this, we expected that the absolute concentration of phycoerythrin would increase when *R. salina* grew in a green-dominated environment. Our results supported this prediction. Because the cryptophyte phycoerythrin is the major light-harvesting pigment in *R. salina* and it absorbs green light better than chlorophylls do, this suggests that *R. salina* is maximizing its ability to capture light in green wavelengths. In red light, we saw that as a proportion of total cellular pigments cryptophyte phycoerythrin decreased by 12% compared to the wide-spectrum control; chlorophyll-*a* counterbalanced this with a 12% increase (though this was not statistically significant). This suggests that *R. salina* exhibited an investment tradeoff in pigment composition in red light, perhaps using chlorophyll-*a*, which absorbs red better than phycoerythrin does, as the primary photosynthetic absorption compound. These particular differences of pigment concentrations and ratios in green and red light were the strongest we observed and are consistent with the theory of chromatic acclimation.

Other patterns, however, cannot be explained by chromatic acclimation. For example, phycoerythrin was present in red light and both chlorophylls were present in green light even though these pigments do not efficiently absorb these corresponding wavelengths. It is unclear why these pigments were not degraded in environments where other pigments would be more useful. Second, pigment composition in a blue environment runs counter to expectations from chromatic acclimation. Chlorophylls absorb blue light better than cryptophyte phycoerythrin, yet in our blue environment *R. salina* produced a large amount of phycoerythrin and a modest amount of chlorophylls, exhibiting the lowest percent of chlorophyll relative to the total pigment concentration in blue light compared to all other treatments. Third, *R. salina* produced an unexpectedly large amount of chlorophyll-*a* in green light. While some chlorophyll-*a* is always necessary for photosynthesis (because chlorophyll complexes mediate the transfer of energy from phycoerythrin to photosystem II in cryptophytes; Scholes, *et al*. 2006), the lower amounts in blue and wide-spectrum light suggest an excess is being produced in green light. Even though other aspects of pigments in red and green light support chromatic acclimation, the high chlorophyll-*a* in green light, the high phycoerythrin in red light, and the preferential presence of phycoerythrin in blue light shows that in some circumstances *R. salina* invests in producing pigments poorly suited to the ambient light environment.

After ten mitotic generations of acclimation in constant environments, it is implausible to attribute these ineffectual pigments to persistence from the past. We therefore examined possible relationships between pigment concentrations to evaluate the plausibility that connections between synthesis pathways could explain these observations. Our analysis did not reveal any significant correlations between chlorophyll and phycoerythrin concentrations, so linked biosynthesis pathways also cannot directly explain our patterns of pigment plasticity. However, we did find a suggestive correlation between transcription of a chlorophyll binding protein gene (chlorophyll A-B binding protein) and phycoerythrin concentration, and two genes related to phycoerythrin synthesis (a phycoerythrin lyase and a potential phycoerythrin α-subunit) correlated with chlorophyll concentrations. These genes are strong candidates for further study of how chlorophyll and phycoerythrin synthesis may be related at a molecular level. It is possible that chlorophyll synthesis may be partially linked to phycoerythrin synthesis in *R. salina*, even though this isn’t reflected in the pigment concentrations themselves. While we have an understanding of how the synthesis pathways for the cryptophyte phycobilins and chlorophylls are structured (Hill & Rowan 1989; Gantt 1996; Scholes *et al*. 2006; Dammeyer & Frankenberg-Dinkel 2008; Overkamp *et al*. 2014), we do not know if or how these synthesis pathways may be co-regulated, particularly with respect to light spectrum. We do, however, know that the chlorophyll binding proteins work to pass energy from membrane chlorophylls and accessory pigments to the photoreaction centers, and this energy transfer is much more efficient between membrane chlorophylls, phycobiliproteins, and the reaction centers than it is between the carotenoids and the reaction centers (Rathbone *et al*. 2020; Sibelik *et al*. 2020), which could explain why 1) we see a suggestive correlation between chlorophyll binding proteins and phycoerythrin concentration and 2) why we see unexpected discrepancies in how other accessory pigments respond to the various light spectra.

The surprisingly low concentrations of chlorophylls in blue light point toward another mechanism contributing to pigment spectral plasticity: high-energy light acclimation of chlorophyll via a mechanism other than quantity. The absorption and photosynthetic efficiency of chlorophyll-*a* is influenced by light spectrum, such that the overall quantum yield of photosynthesis (measured by the amount of oxygen produced per quantum absorbed) increases in high-energy wavelengths, i.e., blue light (Yocum & Blinks 1957; Vadiveloo *et al*. 2015). As a result, there may be less chlorophyll-*a* when *R. salina* is grown in this environment because the chlorophyll-*a* itself became more efficient when acclimated to blue light compared to red or green light. This is consistent with Heidenreich and Richardson (2019), who saw a decrease in cellular chlorophyll concentrations when both phycoerythrin- and phycocyanin-containing cryptophytes were shifted from wide-spectrum to blue-spectrum light, suggesting a possible acclimation response similar to those induced by high light intensity.

Light interception by photoprotective pigments may also influence light available for capture by photosynthetic pigments, and thereby influence photosynthetic pigment composition. Cryptophyte photoprotective pigments, alloxanthin and α-carotene, have been shown to respond to high light intensities (Mendes *et al*. 2018; Kana *et al*. 2019), but less is known about how they respond to changes in light color. Even though alloxanthin and α-carotene both absorb blue wavelengths efficiently, we found the photoprotective pigment concentrations to be lowest in blue light. It is possible that, as with chlorophyll-*a*, absorption efficiency changes with spectrum, and fewer pigment molecules are then needed to absorb the same amount of energy in blue light as in longer wavelengths, but we know of no evidence to indicate whether that happens.

### How do cryptophyte phycoerythrin genes respond to light spectrum at the transcript level?

We expected to see an increase in expression of the cryptophyte phycoerythrin α and β subunit genes in green light to drive chromatic acclimation, and we predicted that expression of these genes would show the same pattern as the pigment concentration. Although our estimate of α subunit gene expression was higher in green than blue or red light, this was not statistically significant, and the overall pattern did not match that of the phycoerythrin pigment. This suggests that post-transcriptional regulatory mechanisms are the primary determinants of phycoerythrin concentrations.

Most striking, however, was the expression disparity between the α and β subunits. The α subunit gene was expressed at a much higher level on average (∼1,670 TPM) compared to the β subunit (∼78 TPM) across all four treatments, and expression of the two subunits was not correlated. This is noteworthy since they compose the overall cryptophyte phycoerythrin structure in a 1:1 molar ratio (αα’ββ), and thus we expected them to be expressed at similar levels and to covary (Richardson 2022). This unexpected relationship between α and β subunit expression could be due to the differences in evolutionary history between the two subunit genes. The α subunit is encoded in the nucleus (Apt et al. 1995; Douglas & Penny 1999; Curtis et al. 2012), which originated from a hypothesized cryptophyte heterotrophic ancestor, and the β subunit is found in the chloroplast genome (Douglas & Penny 1999; Khan et al. 2007; Donaher et al. 2009; Harrop et al. 2014; Kim et al. 2015; Kim et al. 2017), originally descended from the red-algal endosymbiont. A combination of nuclear and plastid photosynthesis-encoding genes is common in many photosynthetic organisms, where genes located in the nucleus encoding proteins that must be imported into the chloroplast (Eberhard *et al*. 2008; Keeling, 2013; Ute *et al*. 2013). Once these proteins are in the chloroplast, they are assembled into larger complexes for full functionality (Celedon and Cline 2013). Proteins that must be assembled in the thylakoid luminal space must cross multiple barriers between the nuclear envelope and the chloroplast membranes. It is possible that increased expression of nuclear genes ensures that sufficient peptides make it to their final destination and any excess proteins are degraded (Eberhard *et al*. 2008), though Gould *et al*. (2007) showed that the phycoerythrin α-subunit isolated from the cryptophyte *Guillardia theta* was able to cross the five membranes between the nucleus where it is synthesized and the thylakoid lumen where it is processed. Alternatively, since chloroplast-encoded transcripts are long-lived (Hosler *et al*. 1989), it is possible that the chloroplast-encoded phycoerythrin β subunit does not need to be transcribed at the same level as the nuclear-encoded α subunit for the overall cryptophyte phycoerythrin protein complex to be synthesized and assembled (Hosler *et al*. 1989; Kim *et al*. 1993).

Kieselbach *et al*. (2018) showed that *Guillardia theta,* another phycoerythrin-containing cryptophyte, has 20 α-subunit genes that can be expressed at the protein level for cells grown in wide-spectrum light. This indicates that there may be a wide pool of α-subunit genes in some cryptophytes that are available when new phycobiliproteins need to be created, and that cryptophytes may alter their phycobiliprotein α-subunit structure as needed by expressing different α subunit genes. This has been suggested as an alternative way that cryptophytes may respond to changes in light availiabity in place of chromatic acclimation. If this hypothesis were true, then it is logical to think that the α gene expression may be expected to be higher than the β. Spangler *et al*. (2022), however, showed that other cryptophytes with phycoerythrin, such as *Proteomonas sulcata,* did not modify their phycobiliprotein α-subunits when exposed to blue or green light for two weeks, which is the approximate length of time *R. salina* was left in various spectra in this study. Because of duration of our experiment, it is unlikely that even if there were multiple α subunits available in the *R. salina* genome, that the protein structure would be altered, but more research into the actual phycobiliprotein structural differences in various spectra is needed to determine this for certain.

### How do photosynthesis-related genes respond to light spectrum at the transcript level?

We saw changes in expression of phycoerythrin subunits and chlorophyll binding proteins in the different spectral environments as mentioned above, along with various photosystem I and II encoding proteins. Of the sixty-six transcripts annotated as potential chlorophyll binding proteins in our assembly, only three were significantly differentially expressed. We also had only three transcripts encoding for phycoerythrin subunits and lyases that were significantly differentially expressed in any of our spectral comparisons. Curiously, these genes did not have any consistent pattern (i.e., not all the chlorophyll binding proteins were upregulated in red or blue light; not all of the phycoerythrin subunits were upregulated in green light, etc.). Transcripts encoding various chlorophyll binding proteins were upregulated in blue light, red light, and green light; transcripts encoding different phycoerythrin subunits and lyases were also upregulated in all three light spectra. Determining why this is so will require detailed functional investigation into the individual proteins.

We found that genes for the RuBisCo large subunit (rbcL) were significantly upregulated in red light compared to blue, even though the small subunit (rbcS) was not differentially expressed across any of the comparisons. RuBisCo is directly involved in carbon assimilation during photosynthesis and is considered the ultimate rate-limiting component in carbon fixation. Changes to RuBisCo subunit gene expression leads to a change in RuBisCo protein synthesis and overall carbon fixation (Pichard *et al*. 1993; Pichard *et al*. 1996; Patel and Berry 2008; Kim *et al*. 2014). Generally, RuBisCo activity and protein synthesis is directly related to light intensity, but spectrum has been suggested to influence RuBisCo gene expression and protein synthesis as well. Eskins *et al*. (1991) found that blue light enhanced protein synthesis and activity over red light in soybean plants, but that this effect was diminished with the addition of far-red light. In a study using *Chlorella vulgaris,* Kim *et al*. (2014) found that rbcL expression was higher in blue light compared to red or wide-spectrum light. Our results possibly differ from these previous studies because of differences in the study organisms and their corresponding pigment complements, leading to increased carbon fixation in red light compared to blue light. However, information on carbon fixation and photosynthesis rates would be needed to determine this with more certainty.

Overall, we did not see many significant differences in expression of pigment-related genes across the different treatments, which is somewhat unexpected in comparisons where we saw significant differences in pigment composition. This discrepancy between the gene expression profile and pigment composition may be a result of post-transcriptional and post-translational modification (Rochaix 1992, 1996; Gruissem *et al*. 1993), which has been shown to occur in response to changes in light intensity and spectrum in plants (Deng *et al*. 1989). Also, decreasing the transcription of chloroplast-encoded genes does not always affect the rates of their corresponding protein synthesis, so it could be that the transcription levels for the pigment-related genes do not reflect the actual pigment concentrations for this reason (i.e., the mRNA transcript:translated protein product ratio is not necessarily 1:1 (Kim *et al*. 1993)). We note that we did not measure pigment concentration and gene expression at multiple time points to determine if the concentrations and levels of expression change over the length of exposure to the treatment environments, so we cannot conclude for certain that these discrepancies are a result of post-transcriptional or post-translational regulation or if this discrepancy is instead an acclimation response (i.e., perhaps at the time in which we sampled for gene expression, transcription of the genes was reduced, but the pigment concentrations still remained higher from previous generations since being placed in the new environments).

### What other functions respond to light spectrum at the transcript level?

Last, we ran differential expression analysis of our complete gene set to test for significant differences in expression of non-photosynthetic genes across spectra. One particularly interesting gene encodes for a domain of an “algal minus-dominance protein,” which is a protein related to the differentiation of mating types in other algal taxa. This gene was upregulated in blue light compared to all other spectral environments in our experiment. In the green alga *Chlamydomonas reinhardtii,* this gene is involved in gametic differentiation and can be regulated by a combination of nitrogen starvation and blue wavelengths, where nitrogen starvation initiates gametogenesis and blue light triggers the completion of the process (Beck and Haring 1996; Ferris and Goodenough 1997; Lin and Goodenough 2007; Chardin *et al*. 2014). Additionally, our annotation pipeline identified 56 genes (though they were not differentially expressed with light spectra) with putative functions similar to the RWP-RK domain-containing transcription factors, which have been shown to be involved in regulating cell differentiation, sexual reproduction, and nitrogen responses in vascular plants, slime molds, and green algae (Konishi & Yanagisawa 2013; Chardin *et al*. 2014; Tedeschi *et al*. 2016; Ota *et al*. 2019). Cryptophytes reproduce asexually, but there has been speculation that sexual reproduction is possible due to observations of cellular fusions (Hoef-Emden & Archibald 2016; Kugrens & Lee 1988) and accounts of dimorphism in clonal cultures (Hill & Wetherbee 1986; Hoef-Emden & Melkonian 2003).

Changes in cell signaling due to shifts in light spectra is a mechanism that has been shown to trigger reproductive switches in other algal groups (Dring 1987; Hoham *et al*. 1997; Hoham *et al*. 2000; Tardu *et al*. 2016) and may be a potential avenue of investigation for understanding switches between reproductive mechanisms in cryptophytes.

We also saw a change in glyceraldehyde 3-phosphate dehydrogenase (GAPDH) expression, which was upregulated in red light compared to green. GAPDH catalyzes the sixth step of glycolysis, though it can also be involved in mRNA regulation, tRNA export, and DNA replication or repair (Huang *et al*. 1989; Sirover 1998; Qiu *et al*. 2020). This shift in GAPDH expression may be indicative of a shift in carbon metabolism or trophic strategy, such as decreasing photosynthetic function in favor of heterotrophic function (i.e., performing mixotrophy). Mixotrophy has been suggested as a form of metabolic function in cryptophytes (Kugrens & Lee 1990; Gervais 1997; Roberts & Laybourn-Parry 1999; Yoo et al. 2017), including in *R. salina* (Lewitus et al. 1991), though there would need to be more experimentation to investigate this in greater detail. Generally, trophic strategy modification and switches in carbon metabolism have been extensively studied with regard to light *intensity* (e.g. Lewitus *et al*. 1991; Caron *et al*. 1993; Rottberger *et al*. 2013; McKie-Krisberg *et al*. 2015), but there are fewer studies on the effects of light *spectrum* on carbon metabolism (e.g., Hamada *et al*. 2003; Das *et al*. 2011).

Many studies examining the physiological effects of red and blue spectral habitats exist (e.g. Figueroa *et al*. 1994; Hoham *et al*. 1997; Ullrich *et al*. 1998; Aguilera *et al*. 2000; Korbee *et al*. 2005; Kim *et al*. 2019), and gene expression work has increased over the years, including work with green algae (Hermsmeier *et al*. 1991; Lee *et al*. 2018; Li *et al*. 2020), brown algae (Deng *et al*. 2012; Wang *et al*. 2013), red algae (Lopez-Figueroa 1991; Tardu *et al*. 2016), and stramenopiles (Takahashi *et al*. 2007; Losi & Gartner 2008). In many of these gene expression studies, blue wavelengths result in a greater number of upregulated genes than red wavelengths, which we also observed in *R. salina*. In all our comparisons, the higher-energy wavelengths of each pairing (blue light in the blue vs. red; green light in the green vs. red; blue light in the blue vs. green) resulted in the greatest number of significantly upregulated genes, regardless of whether the genes were related to photosynthetic function or not. We are unsure if this is a result of high-energy acclimation or if this is simply because higher-energy light triggers a broader net of molecular pathways (potential photomorphogenesis, photoprotection, DNA repair, pigment biosynthesis, etc.) than the lower-energy wavelengths.

Additionally, blue light has been shown to upregulate genes involved in pigment biosynthesis, circadian rhythm, photoreactivation (DNA repair after exposure to UV-B light), regulation of reactive oxygenic species (ROS) during photosynthesis, and photomorphogenesis (growth and reproductive characteristics) (Wang *et al*. 2013; Tardu et al. 2016). Genes upregulated in red light commonly include genes involved in light-harvesting proteins and general photosynthetic function (Wang *et al*. 2013; Deng *et al*. 2012; Lee *et al*. 2018; Tardu *et al*. 2016; Losi & Gartner 2008; Takahashi *et al*. 2007; Hermsmeier *et al*. 1991; Li *et al*. 2020), which is consistent with what we have found in the present study. The potential effects of green light on algal transcriptomes compared to other light spectra is still a widely unexplored avenue of study, given that most of the existing work focuses on the physiological effects of green-light dominated habitats on different algal species and does not include the molecular consequences (Lopez-Figueroa 1991; Hoham *et al*. 1997; Heidenreich & Richardson 2019).

## Conclusion

We quantified differences in pigment composition and gene expression by *R. salina* cultures grown in wide-spectrum, blue-, green-, and red-light environments. *R. salina* appeared to maximize its capacity to capture available wavelengths using its main light-harvesting pigment (cryptophyte phycoerythrin), but the other pigments exhibited more complex responses to light spectrum than can be predicted by the theory of chromatic acclimation. Additionally, cryptophyte phycoerythrin concentrations and expression of phycoerythrin genes are not directly correlated as we expected, and differences observed may be explained by the evolutionary origins of the subunits and by post-transcriptional regulation. We hypothesize that post-transcriptional and post-translational regulatory mechanisms are responsible for discrepancies observed between broader photosynthetic-related gene expression and physiological results.

Overall, photosynthetic-related gene expression is not sensitive to light spectrum, while some non-photosynthetic genes are regulated by light color, which was unexpected. Of particular interest, we have found genes related to sexual reproduction in the *R. salina* transcriptome that could be investigated further. Sexual reproduction in cryptophytes is not well understood and has only been discussed in a handful of studies; thus, future work can use these genes to examine potential triggers of sexual reproduction and to further our understanding of this process in the cryptophyte group.

The mechanisms controlling photosynthetic gene expression and protein synthesis in *R. salina*, and more broadly in the phylum Cryptophyta overall, remains poorly understood.

Because cryptophytes exhibit such a wide diversity of phycobilin types that are only partially associated with phylogenetic history, questions also remain concerning how cryptophytes with different pigment complements, and thus potentially different ecological niches, may respond to shifts in spectral habitat.

## Experimental Procedures

### Growth and Treatment Conditions

#### Baseline cultures

We grew five replicate cultures of *Rhodomonas salina* CCMP 1319 (from the National Center for Marine Algae and Microbiota at the Bigelow Laboratory for Ocean Sciences) in 150 mL of L1-Si media (Guillard & Ryther 1962) in a Conviron walk-in incubator (Controlled Environments, Inc., Manitoba, Canada) kept at 20°C. These replicate cultures were grown under a wide-spectrum light environment (LumiBar Pro LED Light strip, LU50001; LumiGrow, Emeryville, CA, USA) at ∼30 µmol photons m^-2^ s^-1^ on a 12:12 hr light:dark cycle.

Gas exchange and pH was not monitored, but all cultures were grown in the same experimental chamber and in the same media, so these conditions should not have varied. We swirled each replicate culture by hand daily to prevent settling and help aeration.

#### Experimental populations

Once the baseline cultures reached mid-exponential phase (5- 7 days after inoculation), we used the five replicate cultures to inoculate four experimental populations from each by transferring 5 mL of the culture into 300 mL of fresh media for a total of 20 experimental populations. One experimental population derived from each replicate baseline culture was randomly assigned to each light spectrum treatment.

#### Treatment Conditions

We placed all experimental populations in four separate light environments: wide-spectrum; blue-dominated, green-dominated, and red-dominated, each maintained at ∼30 µmol photons m^-2^ s^-1^ at 20°C, which is comparable to low-light conditions cryptophytes usually inhabit in nature. Like the wide-spectrum environment, the blue and red lights were maintained by LumiBar Pro LED light strips (LU50001; LumiGrow, Emeryville, CA, USA), but the green light environment was provided by an EvenGlow® RGB LED panel (Super Bright LEDs Inc., St. Louis, MO, USA). (Spectra for each environment can be found in Supplemental Figure 1.) We left each population to acclimate in each treatment environment for 10 generations (assuming population intrinsic growth rates of 0.39, 0.43, 0.44, and 0.52, per day in wide-spectrum, green, red, and blue light respectively, as quantified by Heidenreich and Richardson 2019 in the same experimental chambers, meaning that the length of time to reach 10 generations for each treatment varied with growth rate). 10 generations is generally accepted as the minimum requirement for algal species to reach balanced growth conditions in new environments (Parkhill *et al*. 2001). During acclimation, we transferred the populations to new media after ∼5 generations (∼7 days) to ensure they remained in nutrient-replete conditions.

After the populations were acclimated, we sampled for pigments and RNA.

### Pigment Analyses

#### Cryptophyte Phycoerythrin Analysis

We calculated cryptophyte phycoerythrin concentrations using the freeze-thaw centrifugation method of Lawrenz *et al*. (2011). We took 15 mL aliquots of each experimental population and centrifuged them at 2,054 g in a Sorvall RC-4B centrifuge for 10 minutes. The supernatant was removed, and the cell pellet was resuspended in 0.1 M phosphate buffer (pH = 6). We then froze the samples at -20°C for a minimum of 24 hours. After freezing, we thawed the samples at 5°C for 24 hours. The thawed samples were then centrifuged at 11,000 g in a Beckman Coulter 18 Microfuge for 5 minutes to remove excess cell material. We measured the absorbance of the remaining supernatant against a phosphate buffer blank in a 1 cm quartz glass cuvette using a Shimadzu UV-VIS 2450 dual-beam spectrophotometer from 400 to 750 nm in 1 nm intervals. Data were scatter-corrected by subtracting the absorbance at 750 nm from the maximum absorption peak (Lawrenz *et al*. 2011). Information on how the concentrations were calculated can be found in the supplemental information.

#### Non-Phycoerythrin Analyses

For determination of chlorophyll-*a*, chlorophyll-*c_2_*, alloxanthin, and α-carotene concentrations, we filtered 5 mL of each experimental population onto a 25 mm Whatman GF/C filter (GE LifeSciences, Buckinghamshire, UK). These samples were then processed using high performance liquid chromatography (HPLC) to obtain pigment concentrations for each sample. Details for analyzing phytoplankton pigment samples using HPLC can be found in Pinckney *et al*. (1996). Filters were freeze-dried overnight, then pigments were extracted for 24 h at -20°C with 750µL of 90% acetone with 50µL of a synthetic carotenoid as an internal standard. The extracted solution was filtered through a 0.45 µm syringe filter, and 250 µL was injected into a Shimadzu HPLC. Chromatograms were analyzed by comparing retention times and absorption spectra to known standards (HDI, Horsholm, Denmark).

Phycobiliprotein concentration cannot be measured with HPLC, which is why the phycoerythrin and non-phycoerythrin pigment concentration measurements were conducted with different methods.

### Statistical Analyses

We first checked the normality and homogeneity of our data using Shapiro-Wilk and Levene’s tests. Then, we ran an Analysis of Variance (ANOVA) with a Tukey post-hoc comparison or a Kruskal-Wallis test with a Dunn’s Multiple Comparison of Means post-hoc comparison to test for significant differences in pigment concentrations across all four treatments. An ANOVA was used for normally-distributed data (phycoerythrin), while the Kruskal-Wallis test was used if data were non-normally distributed (chlorophyll-*a,* chlorophyll- *c_2_*, alloxanthin, and α-carotene).

### RNA Extractions, Sequencing, and Transcriptome Assembly

After taking samples for pigment analyses, we spun the remainder of each culture (270 mL) in 500 mL centrifuge bottles at 3024g for 30 minutes (Beckman Coulter J2-21 centrifuge; JA-20 rotor) to pellet the cell material. The supernatant was removed, and the remaining cell pellet was split into two pre-weighed 2 mL microcentrifuge tubes. We split each pelleted culture into two samples: 1) to ensure we had at least one sample for sequencing if the RNA from one extraction was of poor quality and 2) to allow us to send off technical replicates for sequencing to test for variation within biological replicates.

We spun the microcentrifuge tubes at 3000g (Beckman Coulter 18 Microfuge) for 12 minutes. The supernatant was removed, and then the pellets were weighed. If the pellet mass was between 50-100 mg, we lysed the cells with 1 mL of Bio-Rad PureZOL reagent; if the mass was less than 50 mg, we used 0.5 mL of reagent. The remainder of our extraction protocol followed the standard TRIzol RNA isolation procedure (detailed in the ThermoFisher Scientific Invitrogen TRIzol Reagent User Guide, Pub. No. MAN0001271, Rev. B.0), with the exception of adding a second ethanol wash step prior to elution to increase the purity of the RNA.

We sequenced each sample with 150bp paired-end Illumina sequencing, generating an average of 30,892,204 reads per sample after trimming. Library preparation was performed at Duke using the Illumina Tru-seq RNA poly-A tail enrichment sample library prep kit. We built transcriptome assemblies with Trinity (Grabherr, et al 2011) and Velvet-Oases (Schulz, et al 2012), and combined them with EviGene (Gilbert 2013). Further quality control, RNA-sequencing details and specifications, and transcriptome assembly and annotation protocols are detailed in the Supplemental Material.

### Building the a priori list of photosynthesis-related genes

Prior to differential gene expression analysis, we compiled a list of 99 photosynthesis-related genes known to exist within cryptophyte genomes (Douglas and Penny 1998; Jarvis and Soll 2001; Gould *et al*. 2007; Koziol *et al*. 2007; Khan *et al*. 2007; Overkamp 2014; Takaichi 2011; Neilson *et al*. 2017). These included genes related to pigment synthesis (e.g., the phycoerythrin subunits, chlorophyll-binding proteins), photosystem assembly (e.g., photosystem I and photosystem II proteins), energy synthesis and use (e.g., ATP synthase subunits, cytochrome b6-f complex subunits), the dark reactions (e.g., RuBisCo subunits, light-independent reductases), and helper proteins (e.g., translocons, transport proteins) (Supplemental Table 1). Details of how we obtained the DNA sequences for these genes of interest can be found in the Supplemental Material.

### Differential Gene Expression Analysis

We mapped trimmed paired-end reads back to the final transcriptome assembly and obtained read count data using kallisto (Bray *et al*. 2016). We first examined the distributions of counts across samples and treatments to determine if any samples needed to be removed from the analysis due to batch effects, and to compare data from our technical replicates. We used the technical replication to test for repeatability of the procedures that occurred during sequencing.

There were no significant differences between our technical replicates as indicated by Degust’s quality control measurements (the distribution of counts across samples and treatments, variation in library size, and boxplots of expression levels across technical replicates). Given these results, we dropped the technical replicates from the remaining analyses to keep the dataset approximately the same size for each biological replicate. However, we did end up dropping our biological replicates from n=5 to n=4 because we found one biological replicate that exhibited consistent outliers when using Degust’s quality control measurements and thus dropped that biological replicate from the remaining analysis.

For each spectral comparison (blue vs. red, green vs. red, and blue vs. green), we used Degust (Powell 2019) to run edgeR with a false discovery rate (FDR) corrected *p*-value cutoff of 0.05 and a log fold-change (FC) cutoff of ±2. We ran this analysis at three different scales: first, we tested the expression of cryptophyte phycoerythrin-subunit genes to determine if they were differentially expressed among spectra; second, we tested the *a priori* list of photosynthesis-related genes to investigate photosynthetic pathways more broadly; third, we tested the complete gene set (the total number of contigs expressed in our assembly) to see whether other genes respond to differences in light spectrum. For the photosynthesis gene set, we performed additional analyses to guard against false negatives by relaxing FDR cutoffs to 0.1 and 0.2, as well. Gene expression levels are reported in transcripts per million (TPM).

### Correlation Analysis

We used Pearson’s correlation coefficients (and Spearman’s rank correlation coefficients where data were non-parametric) to assess potential correlations between pigment concentrations and gene expression independent of light treatment (i.e., the correlations detailed below were performed across all 20 experimental populations, not across the 4 light spectra, to capture variation within treatments). We used the corr.test R command with a Holm-Bonferroni adjustment to control for multiple comparisons (part of the “stats” package) for running individual comparisons of interest, which included identifying potential correlations between the following: cryptophyte phycoerythrin pigment concentration and phycoerythrin protein subunits’ transcript expression (both alpha (α) and beta (β) subunits) and cryptophyte phycoerythrin and chlorophyll concentrations. We also did a correlation analysis of the photosynthesis-related genes to all pigment concentrations.

## Acknowledgements

We would like to thank Matthew J. Greenwold for his assistance throughout the transcriptome assembly process, as well as the National Center for Genome Analysis Support at Indiana University and University of South Carolina’s Research Computing center for data storage, computing technology, and resources. We also thank Kristin Heidenreich, Eric Lachenmeyer, and Brady Cunningham for their assistance with growing and maintaining cultures and pigment analyses, Jake Swanson for editorial assistance on this draft, and Savannah Simon and Dylan C. Davis for their help with gene annotation exploration. Last, we thank Jay Pinckney for his guidance in HPLC protocols and statistical analyses, and Yen-Yi Ho and Robert Friedman for their help in training RAS in transcriptomic and bioinformatic procedures. This project was funded by an award from the U. S. National Science Foundation’s Dimensions of Biodiversity program (DEB 1542555) to TLR and JLD. We know of no conflicts of interest for this manuscript.

## Availability of Supporting Data

Raw sequence data has been deposited in the Sequence Read Archive (SRA) under the accession PRJNA749794. This Transcriptome Shotgun Assembly project has been deposited at DDBJ/EMBL/GenBank under the BioProject accession PRJNA749794. The version described in this paper is the first version, PRJNA749794. Pigment and related data have been deposited in Dryad under https://doi.org/10.5061/dryad.8sf7m0cp4.

