## Supplemental Material for "Consequences of light spectra for pigment composition and gene expression in the cryptophyte *Rhodomonas salina*"

*Calculating cryptophyte phycoerythrin concentrations.*

After performing the freeze-thaw centrifugation method to obtain cryptophyte phycobiliprotein samples and absorbance, the concentrations (pg/cell) were calculated using the following calculation:

$$C = \frac{A}{\varepsilon * d}\ x\ MW\ x\ \frac{Vbuffer}{Vsample} x\ \frac{10^{12}}{N}$$

Where $A$ = absorbance of sample, ε = the extinction coefficient for cryptophyte phycoerythrin ($5.67 \times 10^5$ L ·mol$^{-1}$ · cm$^{-1}$; MacColl et al. 1976), $d$ = path length of the cuvette in cm, $MW$ = molecular weight of cryptophyte phycoerythrin (45,000 Da; MacColl *et al.* 1973, 1976), $V_{buffer}$ = volume of buffer in mL, $V_{sample}$ = volume of sample in mL, and $N$ = concentration of cells (cells/L). The $10^{12}$ is the conversion factor to convert the results into pg/cell from g/cell.

*RNA quality control and RNA-sequencing*

We checked the purity of the RNA using a Nanodrop 2000, obtained the concentration with a Qubit 4 Fluorometer, and checked the integrity of the RNA by running a sample of the extracted RNA on a 2% agarose gel at 60V for 1 hr. We considered high quality RNA samples to be those with 260/280 and 260/230 ratios greater than 1.8 and clear rRNA bands observed on the gel with no signs of degradation.

Samples were sent to the Duke University Genome Sequencing Information Manager for 150bp paired-end HiSeq 4000 sequencing targeting 30 million reads per sample. We submitted RNA samples from each of the five biological replicates for each light environment. For three of those biological replicates, we submitted the sample divided into two technical replicates to ascertain technical variation. In total, we obtained sequences from 32 RNA samples [(5 biological replicates + 3 technical replicates) x 4 environments = 32 samples)].

<u>*Transcriptome Assembly and Quality Control*</u>

We checked the quality of the raw reads for each sample with FastQC (Andrews 2010), and then trimmed reads and removed adapter sequences using Trimmomatic (Bolger, *et al.* 2014) (Trimmomatic parameters: ILLUMINACLIP:TreSeq3-PE.fa:2:30:10 HEADCROP:20). After running Trimmomatic, we re-ran FastQC, and any remaining overrepresented adapter sequences observed were manually removed from each sample using cutadapt (Martin 2010). An average of 30,892,204 paired-end reads per sample remained after trimming, and we used these for transcriptome assembly.

We first created a transcriptome of all samples from all light treatments using Trinity (Grabherr, *et al.* 2011). Then, we created several assemblies using kmer lengths of 35, 45, 55, 65, 75, 85, and 95 with Velvet-Oases (Schulz, *et al.* 2012). We merged the separate Velvet-Oases assemblies using the Oases merge function to create one final Velvet-Oases assembly. We then combined the Trinity and Velvet-Oases assemblies with EvidentialGene mRNA transcript assembly software (EviGene; Gilbert 2013) with a kmer length of 75 to obtain a final transcriptome assembly. We used EviGene to correct for the various biases attributed to the different assemblers, which ensured that we were left with a comprehensive and accurate transcriptome assembly.

We removed any remaining rRNA sequences from the final assembly by downloading the rRNA SSU and LSU subunits for *R. salina* from the SILVA database (Quast, *et al.* 2013) and blasting these sequences against our final *R. salina* assembly. Only 101 contigs hit to the rRNA subunits, and these were removed from the final assembly. We then used Benchmarking Universal Single Copy-Orthologs (BUSCO, version 3) to assess the completeness of the transcriptome by searching our assembly against the BUSCO Eukaryota_odb9 (creation date: 02/11/2016), Chlorophyta_odb10 (creation date: 01/12/2017), and Protists_ensembl (creation date: 11/15/2016) datasets (Simao, *et al. 2015*). We used a combination of BUSCO datasets to assess the completeness of our final assembly because *R. salina* does not fit neatly into any specific category. We used the Quality Assessment Tool for Genome Assemblies (QUAST) (Gurevich *et al.* 2013) to obtain descriptive statistics of the transcriptome assembly.

*Gene Annotation*

We used the *de novo* transcriptome annotator dammit (Scott 2018) to annotate our final assembly. This pipeline uses Transdecoder to build gene models and then searches the Pfam-A, Rfam, OrthoDB, and uniref90 protein databases for annotation information with an E-value cutoff of $1x10^{-5}$. The putative transcripts were also run through a gene ontology (GO) pathway analysis using OmicsBox (BioBam Bioinformatics 2019) to obtain functional annotation information.

*Building the a priori list of photosynthesis-related genes (cont.)*

After identifying the 99 photosynthesis-related genes we were interested in from the published literature, we then pulled the nucleotide sequences of these genes from cryptophyte genomes publicly available in the NCBI database. We then used blastn to independently search for these genes in our transcriptome assembly with an E-value cutoff of $1x10^{-5}$. These sequences were used in our differential gene expression analyses to investigate the plasticity of genes related to light capture and photosynthetic function with respect to spectral habitat.

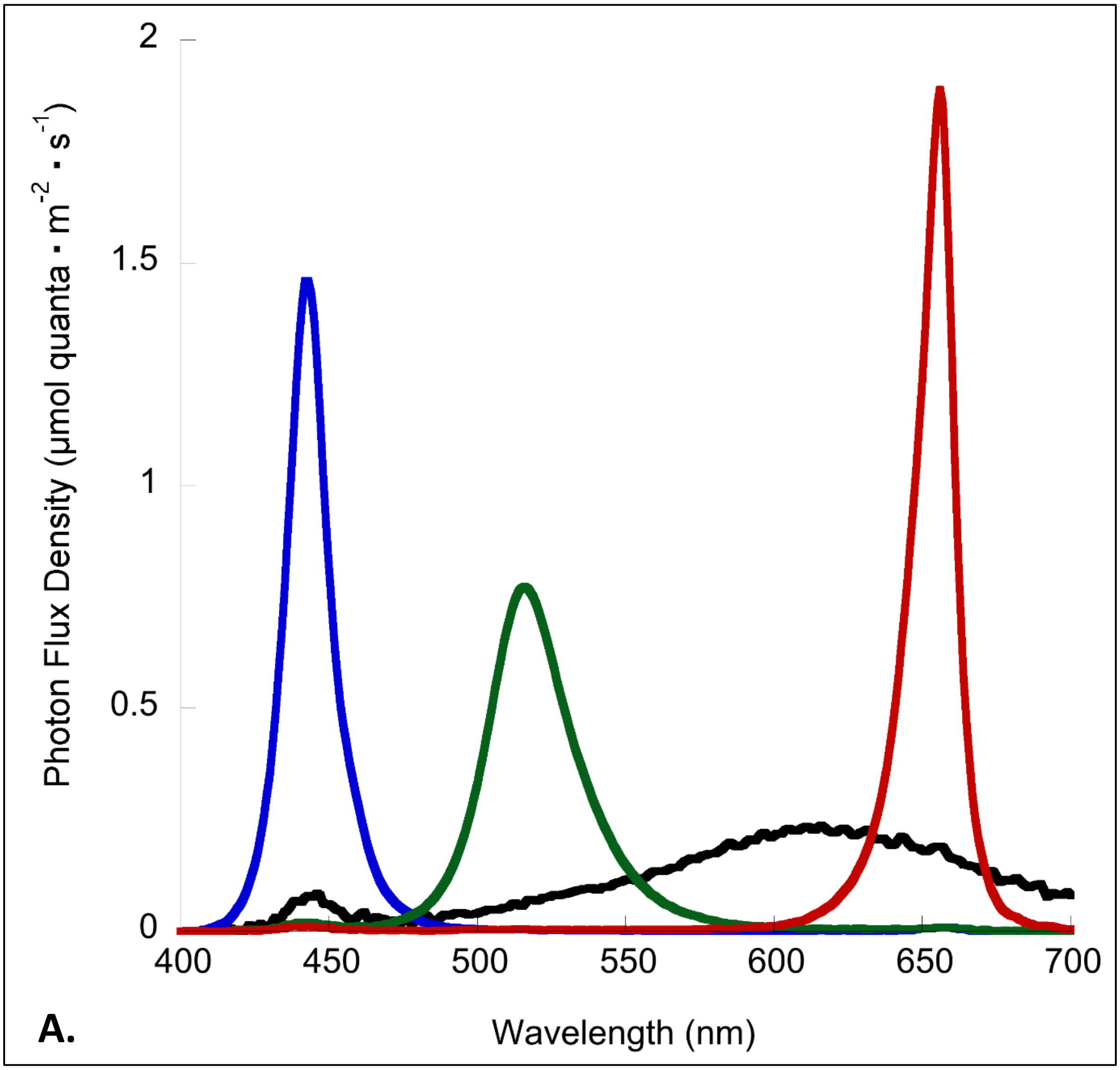

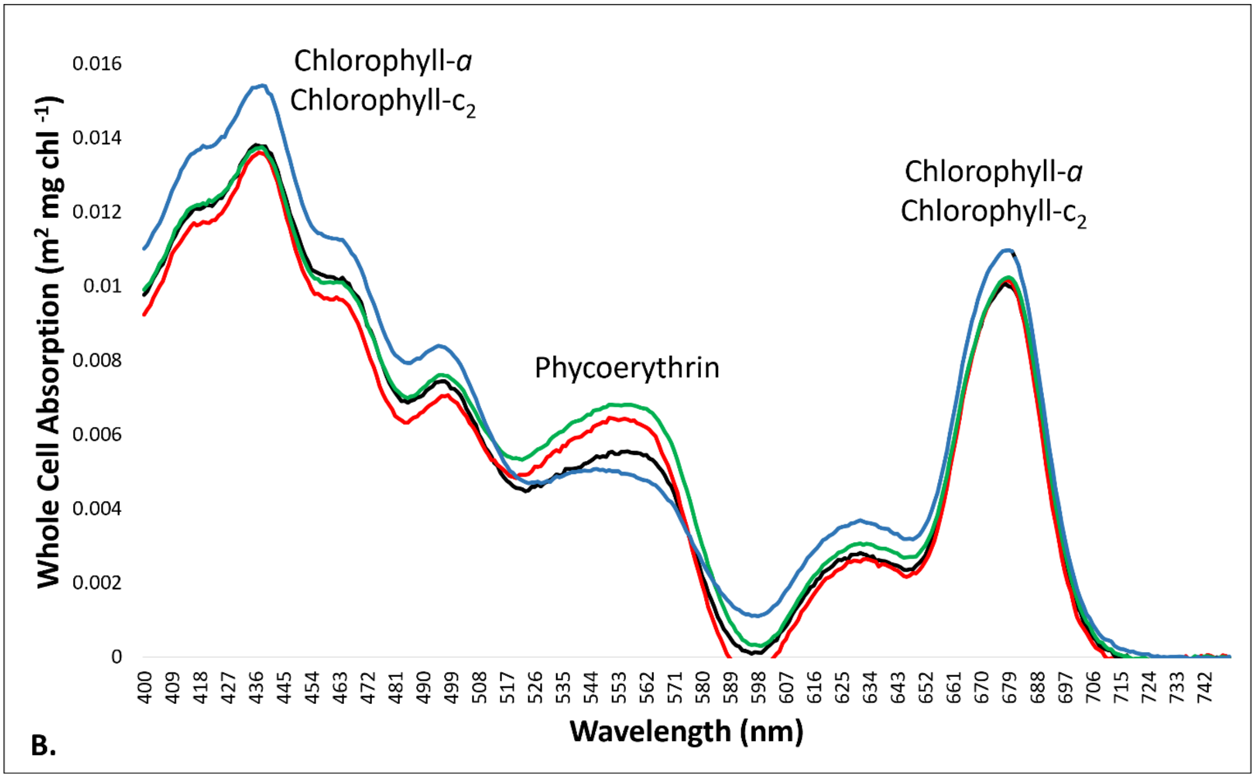


**Supplemental Figure 1:** **A.** Spectral output for each light treatment cultures were grown in. **B.** Whole-cell absorption spectra for *Rhodomonas salina* grown in red, green, blue, and wide-spectrum light. Pigment names are listed near their relevant absorption peaks (Chlorophyll-*a* and chlorophyll-$c^2$ from ~400-500nm and ~620-700nm; Phycoerythrin from ~525-570nm).

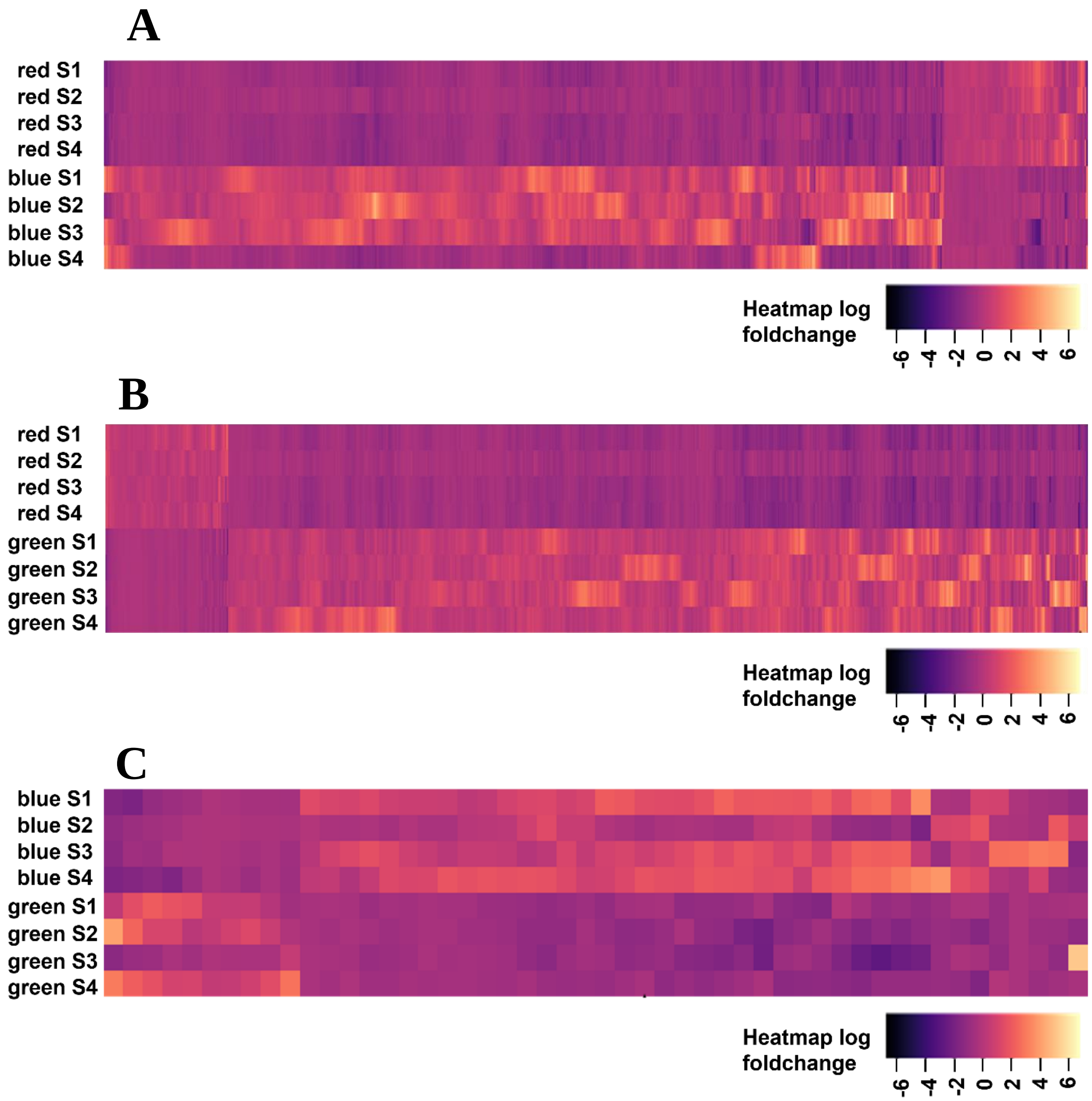


**Supplemental Figure 2:** Heatmaps of the significantly expressed genes for each separate monochromatic comparison. Each heatmap is composed of a different set of genes. **A)** Blue vs. Red; **B)** Green vs. Red; **C)** Blue vs. Green. Each row shows expression from a single replicate population; each column is a different expressed gene. The FDR *p*-value cutoff was 0.05 with a $\log_2$ fold-change > ±2. Upregulated genes are indicated with a positive $\log_2$ foldchange value (yellow); downregulated genes are indicated with a negative $\log_2$ foldchange value (purple).

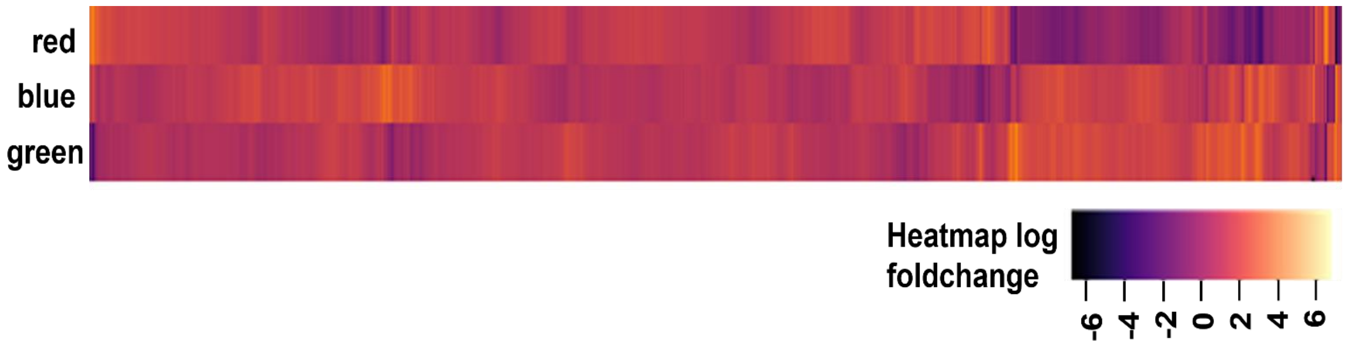


**Supplemental Figure 3** Heatmap of gene expression for all genes across blue, green, and red light environments. This includes all expressed genes in the transcriptome regardless of significance or $log_2$ foldchange value.

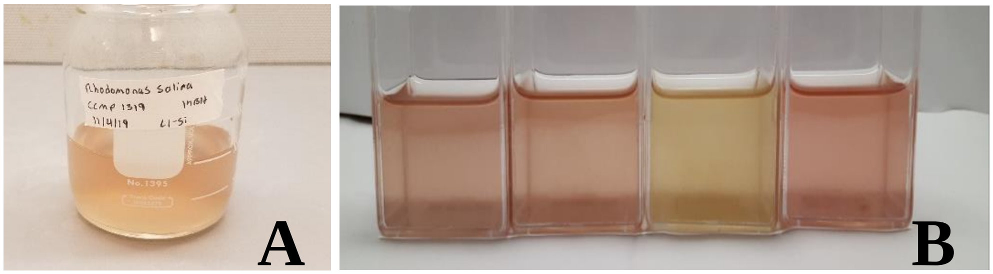


**Supplemental Figure 4: A**) Representative stock culture of *R. salina* grown in wide-spectrum light; **B)** From left to right, *R. salina* cultures grown in wide-spectrum, green, blue, and red light, respectively.

**Supplemental Table 1:** List of the 99 *a priori* photosynthesis-related genes.

| **Gene** | **Symbol** | **GenBank Accession Number** |
|---|---|---|
| Magnesium-Chelatase Subunit, Chloroplastic | chlI | YP_009420404.1 |
| Fucoxanthin Chl A/C Light-Harvesting Protein | lhcr/lhcc | XM_002292317.1 |
| Heme Oxygenase | pbsA | ABO70806.1 |
| Phycoerythrin Alpha Subunit | cpeA | CAC33091.2 |
| Phycoerythrin Beta Subunit | cpeB | ABO70841.1 |
| Phycoerythrin-Associated Linker Genes | cpeC | NC_005125.1 |
| Phycoerythrin-Associated Linker Genes | cpeD | ABP99027.1 |
| Phycobiliprotein/Chromophore Lyase | cpeS | EKX47022.1 |
| Phycoerythrin Alpha Subunit Lyase | cpeZ | CAJ73184.1 |
| Phycoerythrobilin Synthase | pebS | 6QX6_B |
| Phycoerythrin Lyase | cpeT | AFP65495.1 |
| Phycocyanin-Associated Linker Genes | cpcH2I2D2 | AXB27150.1 |
| Phycocyanin-Associated Linker Genes | cpcH3I3D3 | ABQ85421.1 |
| Phycocyanin Alpha Pcb-Like Lyase | cpcX | AIA66941.1 |
| Carotenoid C2-Hydroxylase | crtG | AHB88556.1 |
| Cis-Carotene Isomerase | crtH | QBE68568.1 |
| Lycopene E-Cyclase | crtI | ADD79328.1 |
| Leucopene Cyclase | crtL | ABW31007.1 |
| Phytoene Desaturase | crtP | KAF0161346.1 |
| Carotene Desaturase | crtQ | CAE7265831.1 |
| B-Carotene Hydroxylase | crtR | AUC60050.1 |
| Light Harvesting Apoproteins | LHCP | CAB1097287.1 |
| Light Harvesting Complex | Lhcx | QIH55649.1 |
| Light Harvesting Complex | Lhcz | CAM33414.1 |
| 15,16-Dihydrobiliverdin:Ferredoxin Oxidoreductase | pebA | 6QX6_B |
| Peb:Ferredoxin Oxidoreductase | pebB | 6QX6_A |
| Light Harvesting Protein 2 | lhcb2 | XP_005790942.1 |
| Light Harvesting Complex Protein 8 | lhcr8 | BBO94014.1 |
| Photosystem II Protein D1 | psbA | ABO70840.1 |
| Photosystem II P680 Chlorophyll A Apoprotein | psbB | YP_001293610.1 |
| Photosystem II Cp43 | psbC | YP_001293491.2 |
| Photosystem II D2 Protein | psbD | YP_001293492.1 |
| Cytochrome B559 Subunit Alpha | psbE | ABO70827.1 |
| Cytochrome B559 Subunit Beta | psbF | YP_001293533.1 |

| | | |
|---|---|---|
| Photosystem II Reaction Center Protein H | psbH | YP_001293614.1 |
| Photosystem II Reaction Center Protein I | psbI | YP_001293539.1 |
| Photosystem II Reaction Center Protein J | psbJ | ALG63667.1 |
| Photosystem II Reaction Center Protein K | psbK | YP_001293572.1 |
| Photosystem II Reaction Center Protein L | psbL | YP_001293532.1 |
| Photosystem II Reaction Center N Protein | psbN(A) | ABO70798.1 |
| Photosystem II Reaction Center N Protein | psbN(B) | NP_050801.1 |
| Photosystem II Reaction Center Protein T | psbT | YP_009420501.1 |
| Cytochrome C-550 | psbV | YP_001293523.1 |
| Photosystem II Reaction Center Protein W | psbW | YP_001293482.1 |
| Photosystem II Reaction Center X Protein | psbX | YP_001293524.1 |
| Photosystem II Protein Y | psbY | YP_001293497.1 |
| Photosystem I P700 A Apoprotein A1 | psaA | YP_001293576.1 |
| Photosystem I P700 A Apoprotein A2 | psaB | NC_009573.1 |
| Photosystem I Iron-Sulfur Center | psaC | YP_001293507.1 |
| Photosystem I Subunit II | psaD | YP_001293570.1 |
| Photosystem I Reaction Center Subunit Iv | psaE | YP_001293615.1 |
| Photosystem I Subunit IIi | psaF | YP_001293526.1 |
| Photosystem I Reaction Center Subunit VIIi | psaI | YP_001293530.1 |
| Photosystem I Subunit Ix | psaJ | ALG63666.1 |
| Photosystem I Reaction Center Subunit Psak Precursor | psaK | YP_009420251.1 |
| Photosystem I Reaction Center Subunit Xi | psaL | ALG63562.1 |
| Photosysystem I Reaction Center Subunit XII | psaM | NP_050701.1 |
| Atp Synthase Subunit Alpha | atpA | ALG63566.1 |
| Atp Synthase Subunit A | atpB | NP_050748.1 |
| Atp Synthase Gamma Chain 1 | atpC | NP_050748.1 |
| Atp Synthase Subunit Beta | atpD | YP_001293544.1 |
| Atp Synthase Subunit C | atpE | ALG63580.1 |
| Atp Synthase Subunit B | atpF | YP_001293545.1 |
| Atp Synthase Gamma Chain | atpG | YP_001293546.1 |
| Atp Synthase Cf0 C Subunit | atpH | YP_001293547.1 |
| Atp Synthase Protein I | atpI | YP_001293548.1 |
| Ferredoxin Thioreductase, Beta Subunit | ftrB | YP_001293529.1 |
| Cytochrome F Precursor | petA | YP_001293562.1 |
| Cytochrome B6 | petB | YP_001293480.1 |
| Cytochrome B6/F Compex Subunit 4 | petD | ABO70864.1 |
| Ferredoxin, Chloroplastic Precursor | petF | YP_001293609.1 |
| Cytochrome B6-F Complex Subunit 5 | petG | ABO70761.1 |
| Cytochrome B6-F Complex Subunit 6 | petL | NP_050725.1 |
| Cytochrome B6-F Complex Subunit 7 | petM | ABO70756.1 |

| | | |
|---|---|---|
| Cytochrome B6-F Complex Subunit 8 | petN | NP_050755.1 |
| Cytochrome B559 Subunit Alpha | pbsE | WP_011124483.1 |
| Cytochrome B559 Subunit Beta | pbsF | QXE46209.1 |
| Cytochrome C-550 Precursor | pbsV | WP_011057125.1 |
| Cytoplasmic Polyadenylation Element Binding Protein 1 | cpeB1 | JAO39450.1 |
| Putative Septum Site-Determining Protein Mind Homolog, Chloroplastic | minD | YP_001293500.1 |
| Cell Division Topological Specificity Factor Homolog, Chloroplastic | minE | ABO70851.1 |
| Atp-Dependent Zinc Metalloprotease Ftsh 1, Chloroplastic | ftsH | ABO70796.1 |
| Twin Arginine Transport | TAT | ABO70753.1 |
| Er-Associated Degradation-Like Machinery | ERAD-like | ACS36235.1 |
| Translocon At Inner Membrane Of Chloroplast (Tic) | TIC | KAA8499901.1 |
| Translocon At Outer Membrane Of Chloroplast (Toc) | TOC | XP_002509092.1 |
| Secretory Translocon | SEC | NP_050750.1 |
| Acetate Permease A | acpA | ABO70762.1 |
| Light-Independent Protochlorophyllide Reductase Subunit B | chlB | EU233748.1 |
| Light-Independent Protochlorophyllide Reductase Iron-Sulfur Atp-Binding Protein | chlL | EU233749.1 |
| Light-Independent Protochlorophyllide Reductase Subunit N | chlN | EU233756.1 |
| Acetolactate Synthase Isozyme 1 Large Subunit | ilvB | ABO70722.1 |
| Acetolactate Synthase Isozyme 3 Small Subunit | ilvH | ABO70838.1 |
| Nad-Dependent Dihydropyrimidine Dehydrogenase Subunit Prea | preA | YP_009497974.1 |
| Anthranilate Synthase Component 2 | trpG | YP_009498064.1 |
| Rubisco (Ribulose 1,5 Bisphosphate Carboxylase/Oxygenase), Large Chain | rbcL | ABO70740.1 |
| Rubisco, Small Chain | rbcS | ABO70741.1 |
| High Light Inducible Protein | hlip | ABO70729.1 |
| Chlorophyll binding protein | cab | NP_050676.1 |

**Supplemental Table 2:** Significant correlations between photosynthesis-related genes and chlorophyll-*a* or chlorophyll-$c_2$ pigment concentrations.

| **Gene** | **Chl-*a* *p*-value** | **Chl-*a* Correlation Coefficient** | | **Gene** | **Chl-$c_2$ *p*-value** | **Chl-$c_2$ Correlation Coefficient** |
|---|---|---|---|---|---|---|
| atpH | 0.009 | 0.150 | | CpeT | 0.000 | -0.892 |
| psbV | 0.009 | 0.032 | | Chl_bind32 | 0.012 | -0.613 |
| psbY | 0.024 | -0.116 | | psbW | 0.035 | -0.529 |
| chlI | 0.042 | -0.055 | | PE_ab2 | 0.037 | 0.523 |
| psaA | 0.043 | 0.096 | | Chl_bind19 | 0.043 | 0.510 |
| atpG | 0.044 | -0.207 | | Chl_bind8 | 0.048 | -0.502 |
| atpF | 0.044 | -0.207 | | | | |
| psbH | 0.045 | -0.115 | | | | |

**Supplement Table 3**: Significant correlations between photosynthesis-related genes and photoprotective pigment concentrations.

| **Gene** | **α-carotene *p*-value** | **α-carotene Correlation Coefficent** | | **Gene** | **Alloxanthin *p*-value** | **Alloxanthin Correlation Coeffcient** |
|---|---|---|---|---|---|---|
| atpA | 0.002 | 0.709 | | psbV | 0.008 | 0.634 |
| psbB | 0.003 | 0.691 | | atpH | 0.01 | 0.622 |
| petF | 0.003 | 0.684 | | psbY | 0.038 | -0.523 |
| petG | 0.005 | 0.667 | | chlI | 0.042 | 0.514 |
| atpH | 0.006 | 0.650 | | psaA | 0.048 | 0.501 |
| rbcL | 0.007 | 0.648 | | | | |
| atpB | 0.010 | 0.623 | | | | |
| Chl_bind33 | 0.016 | -0.590 | | | | |
| psbK | 0.017 | 0.586 | | | | |
| psbD | 0.018 | 0.584 | | | | |
| psbC | 0.018 | 0.584 | | | | |
| Chl_bind3 | 0.023 | -0.564 | | | | |
| Chl_bind11 | 0.023 | -0.562 | | | | |
| Chl_bind27 | 0.035 | -0.529 | | | | |
| Chl_bind21 | 0.039 | -0.521 | | | | |
| atpI | 0.039 | 0.521 | | | | |
| PSII.Component1 | 0.041 | -0.515 | | | | |
| Chl_bind30 | 0.046 | -0.506 | | | | |
| psbA | 0.047 | 0.503 | | | | |
